# Molidustat Targets a Synthetic Lethal Vulnerability in APC-Mutant Colorectal Cancer through GSTP1 and PHD2 Co-Inhibition

**DOI:** 10.64898/2026.01.31.702998

**Authors:** Chiara Asselborn, Agata N. Makar, Jair G. Marques, Aslihan B. Akan, Athanasia Yiapanas, Carrie Jennings, Ana Perez Lopez, Jimi Wills, Asier Unciti Broceta, Kevin B. Myant, Alexander von Kriegsheim

## Abstract

Mutations in the adenomatous polyposis coli (APC) gene are a defining feature of colorectal cancer (CRC) and impose metabolic and stress-adaptation requirements that may create exploitable vulnerabilities. Prolyl hydroxylase domain (PHD) inhibitors have been explored as therapeutic agents in CRC, however, their mechanisms of action and off-target effects remain elusive. Serendipitously, we found that Molidustat, a PHD2 inhibitor, induced cell death in APC mutant CRC cells. Ablation of PHD2 alone did not affect cell viability, suggesting an off-target mechanism. Using thermal proteome profiling and chemical proteomics, we identify glutathione S-transferase P1 (GSTP1) as a previously unrecognised off-target of Molidustat and demonstrate direct inhibition of its enzymatic activity. Genetic ablation of PHD2 alone did not phenocopy the cytotoxic effects of Molidustat, whereas combined loss of PHD2 and GSTP1 induced synergistic proteomic changes associated with cell-cycle suppression and apoptotic signalling. Integrated proteomic and metabolomic analyses further revealed energetic and metabolic perturbations specific to simultaneous GSTP1 and PHD2 loss. Consistent with these findings, APC-mutant colonic organoids displayed selective sensitivity to Molidustat that was not reproduced by hydroxylase inhibition alone, supporting a synthetic lethal interaction between GSTP1 and PHD2 in APC-mutant contexts. Together, these results identify a functional interaction between GSTP1 and PHD2 in a subset of colorectal cancer and suggest that off-target engagement of GSTP1 contributes to the anti-tumour activity of Molidustat.

## Introduction

Colorectal cancer (CRC) is the second leading cause of cancer-related death worldwide and a major contributor to global cancer morbidity^1^. Despite advances in screening and targeted therapy, long-term survival remains poor, underscoring the need to identify molecular vulnerabilities that can be exploited therapeutically. Understanding the adaptive mechanisms that enable tumour cells to tolerate oncogenic stress is critical for developing precision approaches to CRC treatment.

Mutations in the adenomatous polyposis coli (APC) gene occur in the vast majority of CRCs and represent the initiating event in tumour development^2,3^. Loss or mutation of APC leads to constitutive activation of Wnt/β-catenin signalling pathway, driving uncontrolled proliferation^4–6^. Moreover, in APC-mutant cells, β-catenin signalling contributes to adaptation programmes that support tumour growth under low oxygen availability (hypoxia)^7^. During hypoxia, oxygen sensing and signal transduction are largely governed by prolyl hydroxylase domain proteins (PHD1/2/3), which hydroxylate target proteins in an oxygen-dependent manner^8,9^. Canonically, PHDs control the stability of hypoxia-inducible factor-1α (HIF-1α) by promoting its proteasomal degradation under normoxic conditions, in which molecular oxygen availability permits efficient hydroxylation and binding to VHL protein^10,11^. However, these hydroxylases also modify non-HIF substrates, and substrate-trapping proteomics has shown that hydroxylase signalling extends to pathways relevant to Wnt regulation^12^. Altogether these observations suggest that modulation of PHD activity may represent an attractive approach for targeting APC-deficient colorectal cancer.

Pharmacological inhibitors of PHDs are widely used to disrupt the hydroxylase function. The most frequently used compound, dimethyloxalylglycine (DMOG), is a pan-PHD inhibitor simultaneously disrupting the function of all 2-Oxogluterate-dependent dioxygenases ^13,14^. In contrast, newer clinical compounds such as Molidustat (BAY 85-3934) were developed with improved selectivity toward distinct PHD enzymes and optimised pharmacological properties^15^. However, selective PHD inhibition can exert broader biological effects: IOX5 has been shown to suppress tumour growth and induce apoptosis in acute myeloid leukaemia through HIF-regulated and off-target mechanisms^16^. Likewise, although Molidustat has been shown to reduce tumour survival in breast cancer models, the mechanism of action and off-targets of this compound remain unknown^17^.

Glutathione S-transferase P1 (GSTP1) is a detoxification enzyme that conjugates glutathione to electrophilic substrates and modulates stress-activated signalling pathways^18,19^. GSTP1 is frequently upregulated in CRC and promotes proliferation and invasion through STAT3 activation^20^. Altered GSTP1 expression or methylation has also been associated with tumour progression and therapy resistance in other cancers^21,22^. Given its role in maintaining cellular stability under stress, GSTP1 may counteract the cytotoxic consequences that arise when oxygen-sensing pathways are perturbed. Consequently, GSTP1 inhibitors are an active area of research, with several compounds at the pre-clinical and clinical stage^23–25^.

Driven by the connection between hydroxylase activity, APC-relevant signalling pathways, and by observations that selective PHD inhibition can affect tumour cell survival, we investigated how PHD inhibition influences adaptive stress responses in APC-mutant colorectal cancer. In this study, we characterise cellular targets of the clinical PHD inhibitor Molidustat and identify a synthetic lethal interaction between GSTP1 and PHD2 inhibition in APC-mutant colorectal cancer models. Using pharmacological inhibition and CRISPR-mediated gene disruption, we demonstrate that concurrent blockade of GSTP1 and PHD2 synergistically disrupts homeostatic control, inducing proteomic reprogramming, accumulative stress, and apoptotic cell death in APC-deficient cells. This dual vulnerability exposes a therapeutic window for selectively targeting APC-mutant colorectal cancers while sparing normal epithelia.

## Methods

### Cell culture and treatments

Human colorectal cancer cell lines HT29 and RKO were obtained from the American Type Culture Collection (ATCC) and maintained in Dulbecco’s Modified Eagle Medium (DMEM; Sigma-Aldrich, D6429) supplemented with 10% (v/v) fetal bovine serum (FBS; Gibco, 16000044), 1% (v/v) penicillin-streptomycin (Gibco, 15140122), and 2 mM L-glutamine (Sigma-Aldrich, G7513). Cells were cultured at 37 °C in a humidified incubator with 5% CO₂.

For chemical inhibition of prolyl hydroxylases (PHDs), cells were treated with Molidustat (BAY 85-3934; Cayman Chemical, 17744), IOX4 (Tocris Bioscience, 5084), DMOG (dimethyloxalylglycine; Cayman Chemical, 71210), or JNJ-42041935 (Sigma-Aldrich, SML1390). For iron chelation-based HIF stabilisation, deferoxamine mesylate (DFO; Sigma-Aldrich, D9533) was used. For GSTP1 inhibition, Ezatiostat (Cayman Chemical, CAY-16248) was used. All compounds were dissolved in DMSO (Sigma-Aldrich, D2650) to prepare 10 mM stocks and diluted in complete medium to the indicated concentrations. Cells were seeded 24 h prior to treatment to achieve 60-70% confluence and incubated with compounds or vehicle control for the specified durations.

### 3D organoid culture

Murine wild-type and Apc^fl/fl^ organoids were generated as previously described^26^. Apc^fl/fl^ intestinal organoids were derived from Villin-CreERT2;Apc^fl/fl^ mice. Briefly, intestinal crypts were isolated from freshly dissected small intestines, washed in cold phosphate-buffered saline (PBS), and incubated in 2 mM EDTA (Thermo Fisher Scientific, 15575020) for 30 min on ice. Crypts were pelleted (300 × g, 5 min, 4 °C), resuspended in growth-factor-reduced Matrigel (Corning, 356231) or BME (Basement Membrane Extract Matrix, Type 2, Trevigen #3533-010-02), and plated in in 24-well plates. After polymerisation (15 min, 37 °C), organoids were overlaid with advanced DMEM/F12 (Gibco, 12634010) supplemented with 10mM HEPES (Invitrogen, 15630106), 1% (v/v) penicillin-streptomycin (Gibco, 15140122), and 2 mM L-glutamine (Sigma-Aldrich, G7513), N2 (Invitrogen, 17502048) and B27 (Invitrogen, 17504044). Medium was also supplemented with EGF (Peprotech, 315-09-500UG), 1% conditioned medium Noggin (produced in-house) WT organoid medium was further supplemented with 10% conditioned medium R-Spondin (produced in-house).

### In vivo Molidustat treatment

All mouse models were bred and kept in the animal facilities of the University of Edinburgh according to UK Home Office regulations. They were kept in 12h cycles of light/dark and had access to water and standard food ad libitum.

Gene deletion in Lgr5-EGFP-ires-CreER^T2^;Apc^fl/fl^ mice^27^ was induced using intraperitoneal injection (IP) of tamoxifen on two consecutive days (120mg/kg on Day 0 and 80mg/kg on Day 1). After induction with tamoxifen, mice were treated daily with Molidustat (10mg/kg, suspended in 2.5% DMSO in PBS) and compared to vehicle-only controls. Once the animals reached the clinical endpoint, they were sacrificed by cervical dislocation. 2 hours prior, the mice were injected with 200µM BrDU (GE Healthcare, RPN201). For IHC, tissues were first washed with PBS and fixed in 4% paraformaldehyde for 24h at 4°C. Tissues were processed in a Tissue-TEK VIP Infiltration Processor (Sakura). The processed samples were embedded in paraffin wax and 5 μm sections cut on a microtome (Leica). Standard immunohistochemistry and histology techniques were used. A 1/500 dilution was used for the BrdU antibody (BD Biosciences, 347580). Images were taken using the Nanozoomer Digital slide scanner (Hamamatsu) with the NDP.view2 software (Hamamatsu). QuPATH (REFERENCE) was used for image analysis and quantification. For the qRT-PCR experiment, which were used to assess the expression of Epo mRNA in the kidney, SYBR Select Master Mix (Applied Biosystems, 4472920) was used according to the protocol and the reaction was performed in a CFX Connect Real-time System (BioRad) machine. Epo forward: 5’-ACTCTCCTTGCTACTGATTCCT-3’, Epo reverse: 5’-ATCGTGACATTTTCTGCC TCC-3’.

### CRISPR/Cas9-mediated knockout

Gene knockouts were generated either using the Edit-R predesigned crRNA (Dharmacon) system or the Alt-R CRISPR-Cas9 system (Integrated DNA Technologies, IDT). For the Dharmacon system, HT29 and RKO cells were transduced with Lentiviral Cas9 (Addgene: #52962), and selected with blasticidine for at least 3 days. Pre-designed PHD2 (EGLN1) crRNAs (CM-004276-01-0002, CM-004276-02-0002, CM-004276-03-0002; targeting sequences: TACAACCAGCATATGCTACA, GTGGCTGCCGAAGCCGAGCC, GATAAGATCACCTGGATCGA, respectively) or GSTP1 targeting sequences (AA GGGAAATAGACCACGGTGTA, AATACCATCCTGCGTCACCT) .were mixed with tracrRNA (Dharmacon, U-002005-05) for 5 min, incubated with DharmaFECT 2 for 20 min, before transfecting the cells. Knockout efficiency was assessed after 3 days. Edit-R Non-targeting crRNA Control #1 (U-007501-01-05) and Edit-R Lethal crRNA Control #1 (U-006000-01-05) were used as controls. For the Alt-R CRISPR-Cas9 system (Integrated DNA Technologies, IDT) system, crRNAs targeting human or mouse PHD2 (EGLN1) and GSTP1 were designed using the IDT design tool. crRNAs were annealed with tracrRNA (IDT, 1072533) by heating at 95 °C for 5 min and cooling to room temperature. Cas9 nuclease (IDT, 1081058) was incubated with the RNA duplex for 15 min to form ribonucleoprotein (RNP) complexes. Cells were nucleofected using the Amaxa 4D-Nucleofector system (Lonza, V4XC-1032) with the SE Cell Line Kit according to the manufacturer’s protocol. Transfected cells were recovered for 48 h in complete medium before selection. Knockout efficiency was validated by Western blotting.

### Caspase-3/7 activity assay

Apoptotic cell death was quantified using the CellEvent Caspase-3/7 Green Detection Reagent (Invitrogen, C10423). Cells were seeded in 96-well plates and treated with inhibitors or vehicle control. Caspase-3/7 reagent (2 µM or 5 µM final concentration) was added directly to the medium, and plates were imaged using the IncuCyte Live-Cell Analysis System. Caspase activity was quantified using IncuCyte software. In organoids, CellEvent Caspase-3/7 Green Detection Reagent (Invitrogen, C10423) was added directly to the medium (2 µM final concentration) and brightfield and fluorescent images were captured manually. Images were analysed using ImageJ (Fiji) software.

### MTT cell viability assay

Organoids were seeded in 96-well plates and cultured for 2 days before treatment with fresh medium containing PHD inhibitors or vehicle control. After 48 h, the medium was removed and replaced with fresh medium containing a final concentration of 5 μg/ml MTT reagent (Sigma, M2128). After 2 h incubation at 37°C, the MTT medium was removed and 2% sodium dodecyl sulfate (SDS) (in H2O) was added to solubilise the Matrigel. After 2 h, DMSO was added to the wells to solubilise the reduced MTT. After 1 h, the optical density was measured on a microplate reader (Wallac, 1420 Victor2 microplate reader) at 570 nm and 690 nm (background).

### Clonogenicity assay

Organoids were seeded in 24-well plates and cultured for 2 days before treatment with Molidustat or vehicle control. After 24 h, organoids were collected, washed and centrifuged at 300 x g (3 min), followed by dissociation to single cells using TrypLE (Gibco, 12604013) for 20 min (37°) and passed through a 40µm cell strainer. 10,000 single cells were resuspended per 10 μl droplet of fresh BME (Trevigen #3533-010-02). The BME droplets were covered in standard organoid culture medium, supplemented with Y27632 dihydrochloride (ROCK inhibitor, Tocris, 1254). After 3 days, brightfield images of the resulting clones were captured and counted using ImageJ (Fiji) software.

### LGR5 Real-time quantitative polymerase chain reaction (RT-qPCR)

Organoids were cultured as described above for 2 days followed by treatment with Molidustat or vehicle control for 48h. Organoid RNA was extracted using the RNeasy Mini Kit (Qiagen, 74106) and DNase treatment (Invitrogen, AM1906) was performed. cDNA was synthesised using 500 ng -1 µg RNA and the qScript cDNA SuperMix (Quantabio, 95048-100). cDNA was diluted (1/10) prior to RT-qPCR on the CFX Connect Real-Time System (Bio Rad). The reaction was prepared using SYBR Select Master Mix (Applied Biosystems, 4472920) and the following LGR5 oligonucleotides: 5’ GAGTCAACCCAAGCCTTAGTATCC 3’ and 3’ CATGGGACAAATGCAACTGAAG 5’

### Protein extraction and Western blotting

Cells were lysed in RIPA buffer (Thermo Fisher Scientific, 89901) supplemented with protease and phosphatase inhibitors (Roche, 11836170001 and 4906837001). Lysates were cleared by centrifugation (14,000 × g, 15 min, 4 °C) and quantified using the BCA Protein Assay Kit (Pierce, 23225). Equal protein amounts were mixed with 4x NuPage LDS buffer (Thermo Fisher Scientific, NP0007), denatured, and separated by 10% SDS-PAGE. Proteins were transferred to PVDF membranes (Millipore, IPVH00010), blocked in 5% non-fat milk/TBS-T, and incubated overnight with primary antibodies at 4 °C. HRP-conjugated secondary antibodies (Cell Signaling Technology) and either Clarity Western ECL substrate (Bio-Rad, 1705061) or SuperSignal West Femto (Thermo Scientific, 34094) were used for detection on a Bio-Rad ChemiDoc XRS+ Imaging System.

### GSTP1 recombinant protein expression and enzymatic assays

The plasmid used for recombinant expression of human GSTP1 was kindly provided by Dr Anna-Maria Caccuri (University of Rome). Recombinant GSTP1 was expressed in E. coli BL21 (DE3) cells induced with 0.5 mM IPTG (Sigma-Aldrich, I6758) for 4 h at 30 °C. Cells were lysed by sonication in PBS containing 1 mM PMSF (Sigma-Aldrich, P7626), and the fusion protein was purified using Glutathione Sepharose 4B resin (Cytiva, 17-0756-05). GSTP1 was eluted with 10 mM reduced glutathione (Sigma-Aldrich, G4251). Enzymatic activity was measured spectrophotometrically at 340 nm using glutathione (GSH) and 1-chloro-2,4-dinitrobenzene (CDNB; Sigma-Aldrich, 138630) in 100 mM potassium phosphate buffer (pH 6.5). For inhibition assays, purified GSTP1 (0.5 µg) was pre-incubated with ethacrynic acid (Sigma-Aldrich, E8750; 1-100 µM, 30 min, room temperature) or Molidustat (BAY 85-3934; Cayman Chemical, 17744) before substrate addition. Enzymatic activity was calculated as the change in absorbance per minute relative to control reactions.

### In silico docking

The conformational energy of the compounds were minimised using Avogadro v.1.2.0. The compounds were docked using a modified protein structure from the Protein Data Bank (https://www.rcsb.org/structure/5X79) using GOLD software in Hermes through Cambridge Crystallography Database (CCDC) v.2025.1. The highest docking scores were visualised in PyMol v.3.1.5.1, and the figures were generated using PyMol v.3.1.5.1.

### Reactive oxygen species and antioxidant treatment

Reactive oxygen species (ROS) were detected using the Peroxy Orange-1 (PO1) fluorescent probe (Tocris, Cat. No. 4944). Cells were pre-incubated with 5 µM PO1 in phenol-red-free DMEM for 20 min at 37 °C, followed by incubation with fresh phenol-red-free DMEM containing Molidustat. PO1 was measured 90 min after addition of Molidustat using a plate reader (Tecan Spark 20, excitation wavelength 543 nm, emission 545-750 nm). For antioxidant rescue, cells were treated with 5 mM N-acetyl-L-cysteine (NAC; Sigma-Aldrich, A7250) in addition to Molidustat. ROS levels were normalised to untreated controls.

### Synthesis of IOX4 beads

The ester group of IOX4 (Tocris Bioscience, 5084; 25 mg, 0.075 mmol) was cleaved by treatment with 90% (v/v) trifluoroacetic acid in distilled water at room temperature for 2 h, following a previously reported procedure^28^. After generation of the carboxylic group, the solvent was evaporated under reduced pressure, and the resulting carboxyl-derivative of IOX4 was used directly without further purification. ω-Aminohexyl-Sepharose 4B beads, preswollen in 20% ethanol (Sigma-Aldrich, A6011; 1 mL settled gel, amine loading 8 μmol mL⁻¹), were transferred to a fritted solid-phase synthesis vessel equipped with a 20 µm polyethylene frit and washed twice with 1 mL of distilled water. Solutions containing carboxyl-derivative of IOX4 (8, 4, 2, 1, or 0.5 μmol) and N-ethyl-N′-(3-dimethylaminopropyl)carbodiimide hydrochloride (EDC; 80, 40, 20, 10, or 5 μmol, respectively) were prepared in 1 mL of distilled water, maintaining a 1:10 molar ratio of IOX4 derivative to EDC as recommended by the commercial protocol. Each solution was added to the pre-washed beads (1 mL, 8 μmol amine loading), and the suspensions were gently rotated at room temperature overnight to allow amide bond formation. As a control, ω-Aminohexyl-Sepharose 4B beads (1 mL, 8 μmol) were coupled to maleic acid (4 μmol) under identical carbodiimide-mediated coupling conditions. Following coupling, all bead samples were filtered and washed thoroughly with distilled water (≥3×). Qualitative ninhydrin and TNBS assays revealed a progressive decrease in free primary amines with increasing free-carboxy IOX4 equivalents, consistent with the expected coupling stoichiometry. Residual free amino groups on the beads were capped by treatment with acetic anhydride (80 μmol) and N,N-diisopropylethylamine (DIPEA; 200 μmol) in 1 mL of 50% (v/v) dioxane in distilled water. The reaction mixtures were gently rotated at room temperature for 30 min. The beads were then filtered and washed extensively with distilled water (≥3×), and successful capping was confirmed by qualitative ninhydrin and TNBS assays, confirming the generation of IOX4-Sepharose beads^29,30^.

### IOX4 assay

HT29 cells were lysed in 50 mM Tris-HCl (pH 7.4), 5% glycerol, 150 mM NaCl, 1.5 mM MgCl₂, 0.4% NP-40, 200 µM ascorbic acid, and 10 µM FeSO₄ with one EDTA-free protease inhibitor tablet (Roche, 11873580001) per 25 ml. Lysates were treated with benzonase (250 U ml⁻¹; Sigma-Aldrich, E1014) on ice for 1 h, centrifuged (20,000 × g, 1 h, 4 °C), and diluted 1:1 with detergent-free pulldown buffer. Protein concentration was adjusted to 1 mg ml⁻¹ (BCA assay; Thermo Fisher Scientific, 23225). Lysates were pre-cleared with control beads (30 min, 4 °C) and incubated with Molidustat (10 µM) or DMSO (30 min, 4 °C). Equilibrated IOX4-Sepharose beads (35 µl) were then added and incubated for 30 min at 4 °C. For analysis by SDS-PAGE and Western blotting, proteins were eluted (30 min, 50 °C) in 2× NuPAGE LDS sample buffer (Thermo Fisher Scientific, NP0007) containing 50 mM DTT (Sigma-Aldrich, D0632). For proteomics analysis, beads were washed twice in lysis buffer and twice in pulldown buffer, and first eluted in 2M Urea, 50 mM Tris-HCl pH 7.5 and Trypsin (12.5 ng/μl) for 30 min at 27°C. The same beads were then eluted in 2M Urea, 50 mM Tris-HCl and 1 nM DTT. The supernatants from both elutions were combined and trypsinised overnight in a humidified chamber at 37°C. The peptides were alkylated with Iodoacetamide (IAA) for 30 min (room temperature, in the dark), resuspended in trifluoroacetic acid (TFA; Thermo Scientific, 28901) and desalted with C18 stagetips prior to LC-MS/MS analysis.

### Thermal proteome profiling (TPP)

Cells were treated with Molidustat (10 µM, 1 h) or DMSO prior to collection. Aliquots were heated at 37-67 °C for 3 min in a thermal cycler, lysed in 1% NP-40 buffer containing protease inhibitors, and centrifuged (20,000 × g, 20 min) to obtain soluble fractions. To remove any traces of detergent, samples were precipitated several times in ice-cold acetone (overnight, - 20°C). The next day, samples were centrifuged (20,000 × g, 20 min) and washed several times with fresh ice-cold acetone. A bath sonicator was used, to ensure the pellet was fully resuspended between washes. The samples were reduced and alkylated (6 M GuHCl, 0.5 M Tris-HCl pH 8.5, 1 mg ml⁻¹ chloroacetamide1.5 mg ml⁻¹ TCEP), and digested first with LysC (Promega) for 4 h at 37 °C on a shaker (protease-to-protein ratio of 1:200), followed by 1 µg ml⁻¹ MS-grade trypsin (Promega, V5111) digestion overnight, (. Reactions were stopped with trifluoroacetic acid (TFA; Thermo Scientific, 28901). Peptides were desalted on homemade C18 StageTips and resuspended in 50 mM triethylammonium bicarbonate before labelling with Tandem Mass Tags (TMT-10plex; Thermo Fisher Scientific, 90110). Peptides were desalted again on commercial C18 StageTips and fractionated off-line prior to LC-MS/MS analysis.

Thermal stability profiles were generated using the Easy50 pipeline adapted from Savitski et al.^31^. Protein abundance was normalised to the 37 °C reference, and nonlinear least-squares regression was used to fit sigmoidal denaturation curves following the equation: y = 1 / (1 + exp((Tm - x)/b)), where Tm represents the apparent melting temperature and b the slope parameter. Proteins showing reproducible >1 °C shifts in Tm (p < 0.05) and/or reproducible changes (Δ > 0.1) in soluble abundance at 67 °C (p < 0.05) between treatment and control were considered compound interactors. Volcano plots were created using Prism GraphPad.

### Proteomics analyses

Cells were lysed in PAC buffer (5% SDS, 100 mM Tris-HCl pH 8.5, 1 mg ml⁻¹ chloroacetamide, 1.5 mg ml⁻¹ TCEP), heated at 95 °C for 30 min, and sonicated. Lysates were processed for automated digestion (8 h) on the KingFisher Duo (Thermo Fisher Scientific) using MagReSyn HILIC beads (ReSyn Biosciences) and 70% acetonitrile. Washing was performed with 95% acetonitrile and 70% ethanol, and digestion was carried out with 1 µg ml⁻¹ MS-grade trypsin (Promega) in 50 mM triethylammonium bicarbonate. Reactions were stopped with trifluoroacetic acid (TFA; Thermo Scientific, 28901). Peptides were desalted on C18 StageTips and analysed on an Orbitrap Fusion Lumos Tribrid Mass Spectrometer (Thermo Fisher Scientific) coupled to an Ultimate 3000 Nano LC with a C18 packed emitter (Aurora, IonOptiks). Peptides were separated using a 40 min acetonitrile gradient followed by a 10 min 80% wash in 0.5% acetic acid. Data-independent acquisition (DIA) was performed across m/z 350-1650 at 120k resolution, followed by MS/MS on 45 overlapping windows (200-2000 Da) at 30k resolution (NCE 28). Data were analysed using DIA-NN (version 2.3.0 Academia) or FragPipe platform (version 23.1) against the *Homo sapiens* UniProt reference proteome. Statistical calculations, such as Student’s t-tests, ANOVA, and z-scores, were performed using Perseus software^32^. Heatmaps and hierarchical clustering were conducted using the Morpheus online tool (https://software.broadinstitute.org/Morpheus). Volcano plots were created using the VolcaNoseR online tool^33^. Gene ontology analyses were obtained using the ShinyGO 0.80 online tool^34^ and The Molecular Signature Databases (MSigDB)^35,36^. Proteomics synergy analyses input intensities were log2-transformed and normalised by median-centring. For each protein, an ordinary least-squares linear model was fitted to quantify genetic interactions between the two knockouts. The knockout representing the deviation from additivity on the log2 scale and was used as the synergy score. The null hypothesis of no interaction was tested per protein, and p-values were adjusted across proteins using the Benjamini–Hochberg false discovery rate (FDR); unless noted otherwise, significance was defined at FDR ≤ 0.05. To focus on coherent genetic interactions, we reported a subset of proteins in which both single knockouts changed in the same direction relative to Control (increase or decrease) by at least 0.1 log2 units. All analyses were performed in Python using the pandas and NumPy libraries, with statistical modelling conducted in statsmodels (ordinary least squares with HC3 heteroscedasticity-consistent standard errors) and multiple-testing correction applied using the Benjamini–Hochberg false discovery rate procedure.

Raw datasets have been uploaded to ProteomeXchange under the identifier: PXD071906. The dataset is currently private and available for peer review using the following reviewer credentials: Project accession: PXD071906 Token: yFS4RlR3IxWP. Alternatively, reviewer can access the dataset by logging in to the PRIDE website using the following account details: Username, Password: ucgRvRiekZkl. The data will be made publicly accessible upon publication.

### Metabolomics analyses

For LC-MS metabolomics, 5 × 10⁵ cells per well were seeded in 6-well plates and cultured for 24 h. Metabolites were extracted using 50% methanol, 30% acetonitrile, and 20% water (v/v) for 1 h at -20 °C, followed by centrifugation (16,100 × g, 10 min, 4 °C). Supernatants were stored at -75 °C prior to analysis. Samples were randomised and analysed on a Dionex Ultimate 3000 UHPLC (Thermo Fisher Scientific) coupled to a Q Exactive Hybrid Orbitrap MS (Thermo Fisher Scientific). Metabolites were separated by HILIC using a ZIC-pHILIC analytical column (2.1 × 150 mm; SeQuant-MerckMillipore) with a guard column (2.1 × 20 mm). Mobile phase A contained 20 mM ammonium carbonate with 0.01% ammonium hydroxide; mobile phase B was acetonitrile. A linear gradient from 95% to 5% B over 20 min was followed by re-equilibration for 7 min at 200 µl min⁻¹, 45 °C, with a 5 µl injection volume. MS data were acquired in polarity-switching mode (m/z 70-900) at 70,000 FWHM resolution. Metabolites were identified by accurate mass (<5 ppm) and retention time matching to an in-house standard library. Data were processed using MetaboAnalyst 5.0, Morpheus online tool.

## Results

### Molidustat Induces Cell Death in Colonic Cell Lines Independently of PHD Inhibition

Using a substrate trap approach, we previously identified multiple components of the WNT-signalling pathway as interactors of PHD hydroxylases^12^, suggesting that hydroxylase inhibition may modulate WNT signalling. We hypothesised that perturbing WNT signalling beyond the “just-right” window required for colon cancer progression could impair proliferation and/or induce cell death, offering a potential therapeutic strategy. In murine colitis models, hydroxylase inhibitors were well tolerated and reduced inflammation^37,38^, supporting investigation of their effects in canonical, non-inflammation-driven colon cancer. We employed two well-studied colon cancer cell lines: RKO, which lacks mutations in the WNT pathway, and HT29, which carries a canonical APC truncation^39,40^. Cells were treated with increasing concentrations of Molidustat, a clinically approved inhibitor of PHD2, the most abundantly expressed EglN/PHD hydroxylase in these models^10,11^. In HT29 cells, Molidustat induced a clear dose-dependent increase in Caspase-3/7 activity, whereas RKO cells responded only at the highest concentration (Fig. 1A, B). The effect appeared to be independent of HIF stabilisation as Molidustat stabilised HIF1α and induced the expression of its downstream targets, to a similar extent in both cell lines (Supplementary Fig. S1 A, B). To determine whether PHD2 inhibition accounted for this effect, we generated PHD2-knockout HT29 cells using CRISPR-Cas9 (Fig. 1C). Surprisingly, PHD2 loss neither reduced proliferation nor induced Caspase-3/7 activity (Fig. 1D, E). Thus, although Molidustat is characterised as a PHD2 inhibitor, its pro-apoptotic effects cannot be explained by PHD2 inhibition alone. As Molidustat may interact with additional proteins not affected by PHD2 knockout, we hypothesised that the compound exerts the pro-apoptotic function via other targets.

**Figure 1.**
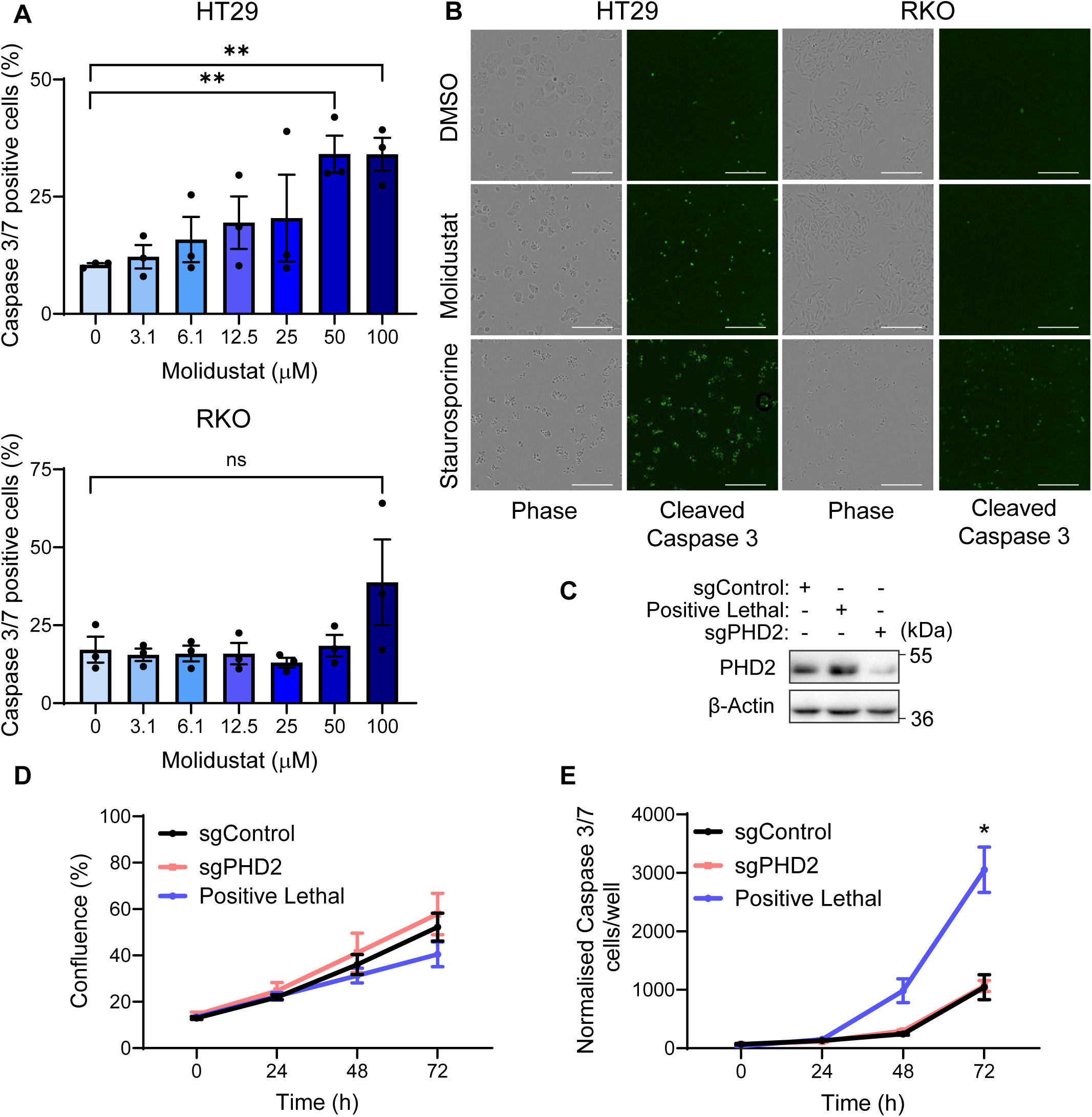
Molidustat Induces Cell Death in Colonic Cell Lines independently of PHD-inhibition. A. Percentage of Cleaved Caspase 3/7 positive HT29 (top), and RKO (bottom) cells. Cells were treated with Molidustat for 48 hours at indicated concentrations, 10uM Staurosporine was used as a positive control (100% cell death). Mean + SEM is assessed by unpaired two tailed Student’s t-test, **p<0.01, (ns) non-significant. B. Representative images of Cleaved Caspase-3/7 signal in DMSO, Molidustat (90 μM), and Staurosporine treated cells. Scale bar: 300 μm. C. Representative Western Blot of PHD2 levels in HT29 cells. D. Percentage confluency of HT29 cells post-transfection with the indicated guide RNAs. E. Cleaved Caspase-3/7 signal in HT29 cells post-transfection with the indicated crRNAs. Mean + SEM is assessed by two-way ANOVA, *p<0.05. *N* = 3 biologically independent experiments.

### Thermal Proteome Profiling Reveals Multiple Molidustat Targets

To identify proteome-wide targets of Molidustat, we performed Thermal Proteome Profiling (TPP)^41^. TPP determines the temperature-dependent denaturation profiles of proteins across the proteome (Fig. 2A), enabling systematic identification of putative drug-protein interactions. HT29 cells were treated in triplicate with Molidustat or DMSO for 1 h, aliquoted, heated between 40-67 °C, lysed, and fractionated to separate denatured from soluble proteins. Soluble fractions were processed and analysed by LC-MS/MS. Melting curves were normalised using empirically defined thermostable proteins and analysed with two independent algorithms. Proteins exhibiting a thermal shift >1 °C (ΔTm50) and/or a significant change in soluble abundance at 67 °C (ΔY) were classified as hits (Fig. 2B, C). A total of 295 proteins met these criteria, including PHD2 (Supplementary Table S1). Consistent with previous reports, Molidustat did not alter the stability of other dioxygenases such as PLOD1 (Supplementary Fig. 1 C-F). Twelve proteins were identified by both algorithms (Fig. 2D). However, altered thermal stability can arise either from direct Molidustat binding or from Molidustat-induced post-translational modifications affecting protein stability^42^. Additional methods are therefore required to distinguish true interactors from indirect effects. Overall, the observed thermal shifts indicate that Molidustat perturbs the stability of a broad set of proteins, consistent with engagement of multiple cellular targets.

**Figure 2.**
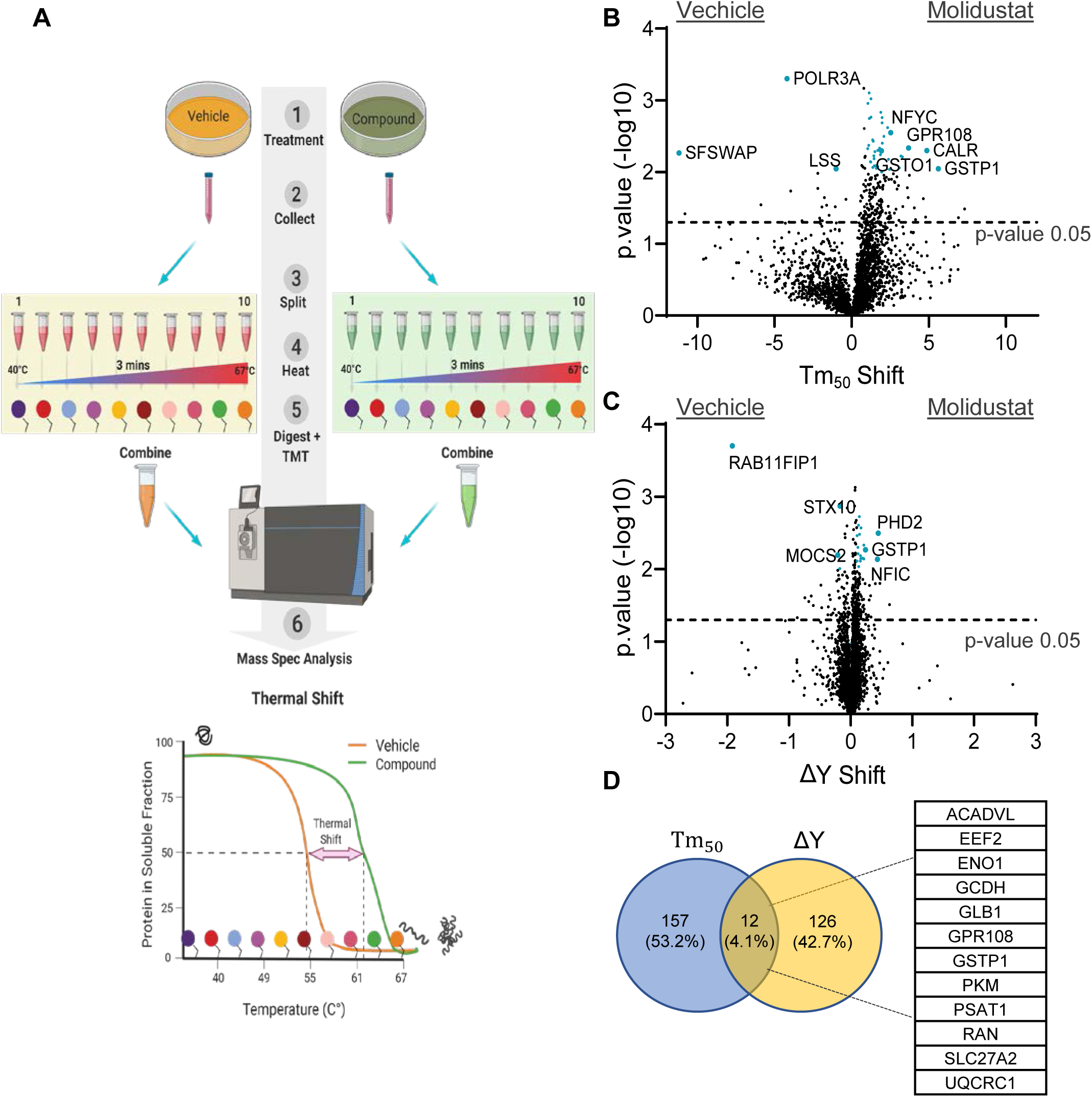
Thermal Proteome Profiling Reveals Multiple Molidustat Targets. A. Intact cells are treated with compound or vehicle control (1). Cells are collected (2), split into fractions (3) with each fraction heated to a different temperature (4). After heat treatment, cells are lysed and proteins are digested to peptides (5). The peptides in each fraction are labelled with a unique TMT tag, which can analysed using mass spectrometer (6), generating melting curves and Tm50 values for the whole proteome. B. Volcano plot of hits identified using the Tm50 method. Tm50 shift and -log10 p-value are shown. Proteins exhibiting a significant shift (< 0.05) greater than 1 °C were considered potential targets. C. Volcano plot of all hits identified using the ΔY method. ΔY shift and -log10 p-value are shown. Proteins with a significant increase (< 0.05) in ΔY greater than 0.1 were taken forward as potential hits. D. Venn diagram showing the overlap of targets identified by both methods. Twelve proteins were identified by both methods are shown on the right.

### Molidustat Inhibits GSTP1 Activity

To refine target identification, we implemented an orthogonal chemical proteomic competition assay resembling an affinity pulldown^43^ (Fig. 3A). Protein lysates were incubated with an immobilised compound or control matrix in the presence or absence of Molidustat. In the absence of free drug, interacting proteins co-precipitated; in its presence, binding sites were blocked, reducing their recovery. Comparing precipitated proteomes with and without Molidustat pre-treatment enabled the identification of specific binders. We generated the affinity matrix by coupling a close analogue of Molidustat IOX4^44^ to agarose beads and optimised coupling conditions using the characterised IOX4-PHD2 interaction (Fig. 3B, C). Mass spectrometry analyses identified multiple proteins altered where pre-treatment with Molidustat reduced the affinity to the IOX4 beads (Supplementary Table S2). Of those, GSTP1 showed the most striking changes in the precipitation levels and Western Blot analyses confirmed that pre-incubation with Molidustat prevented GSTP1 binding to IOX4 beads (Fig. 3D, Supplementary Fig. 2A). GSTP1 also appeared as a hit in the TPP dataset (Fig. 2B, C), supporting its classification as a *bona fide* Molidustat interactor. Consistent with inhibition of GSTP1, a key cellular detoxification enzyme, Molidustat treatment led to a significant increase in intracellular reactive oxygen species (ROS) levels. Co-treatment with the antioxidant N-acetylcysteine (NAC), a ROS scavenger^45^, abolished the Molidustat-induced ROS increase, supporting a functional link between GSTP1 inhibition and redox imbalance (Supplementary Fig. 2B). To further explore the molecular basis of GSTP1 binding, we performed *in silico* docking analyses of Molidustat^46,47,48,49^, which indicated a potential binding site spanning the canonical GST-binding and hydrophobic pocket, which could prevent substrate/GST binding (Supplementary Fig. 2 C). To test whether binding of Molidustat to GSTP1 inhibited in GST-activity, we measured GSTP1 activity using recombinant GSTP1 and the synthetic substrate 1-chloro-2,4-dinitrobenzene (CDNB)^50^. Molidustat inhibited GSTP1 activity in a dose-dependent manner (Fig. 3F), comparable to inhibition by Ethacrynic Acid (EA), another known GSTP1 inhibitor^51^ (Supplementary Fig. 2D). To assess intracellular activity, we repeated the assay in intact HT29 cells and observed reduced GSTP1 activity dose-dependently, whereas EA had no detectable effect (Fig. 3F; Supplementary Fig. 2D). Given the observed increase in ROS, we tested whether pharmacological inhibition of GSTP1 could recapitulate the Molidustat-induced apoptosis in PHD2 knockout cells using Ezatiostat. However, no significant differences in Caspase-3/7 signal were observed between wild-type cells and GSTP1 knockout controls (Supplementary Fig. 2E), rendering these results inconclusive. Taken together, our chemical proteomic, biochemical, and structural analyses identify GSTP1 as a prominent off-target of Molidustat and demonstrate direct inhibition of its enzymatic activity.

**Figure 3.**
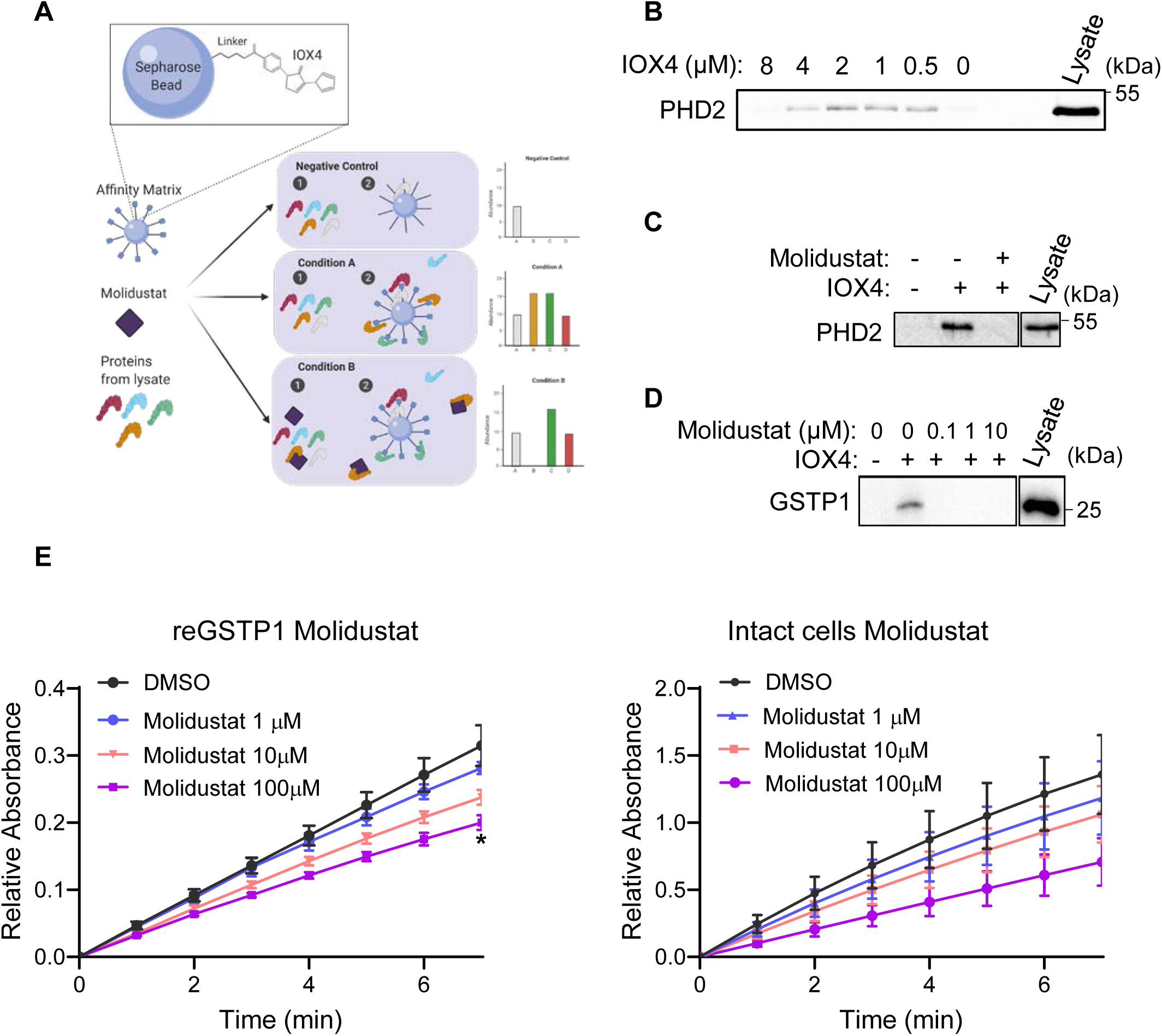
Molidustat Inhibits GSTP1 Activity. A. ω-aminohexyl sepharose beads were coupled with IOX4. Cell lysate is washed over empty beads (negative controls), IOX4 coupled beads (condition A), or IOX4 coupled beads after incubation with Molidustat (condition B). Samples are analysed by Western Blot or Mass Spectrometry. B. Western blot of PHD2 pull-down with IOX4-conjugated beads across a concentration range (0.5-8 μM). C. Competition assay with Molidustat. Lysates were pre-incubated with Molidustat prior to incubation with IOX4-conjugated beads, PHD2 pulldown was assessed by Western Blot. D. Western Blot of GSTP1 binding to IOX4 beads with or without prior incubation with Molidustat. E. Recombinant (left) or cell-based (right) GSTP1 activity assay with Molidustat (1, 10, 100 μM) or DMSO; time-course of CDNB-GSH conjugation monitored at 340 nm and plotted as relative absorbance (arbitrary units, AU). Mean + SEM is assessed by two-way ANOVA, *p<0.05 *N* = 3 biologically independent experiments.

### Combined PHD2 And GSTP1 Loss Drives Synergistic Proteomic Changes Associated With Apoptosis

To test whether genetic ablation of GSTP1 and PHD2 recapitulated the apoptotic phenotype induced by Molidustat (Fig. 1A B), we generated single and double knockout HT29 cell lines and profiled them using quantitative proteomics (Supplementary Table S3). LFQ analysis confirmed efficient loss of GSTP1 and PHD2, and HIF1A accumulation in PHD2-deficient backgrounds (Supplementary Fig. 3A, B). Unsupervised clustering of the top 500 ANOVA-variable proteins identified two major expression clusters, with the most pronounced alterations occurring in the double knockout (Fig. 4A). Notably, loss of either PHD2 or GSTP1 alone modulated a shared subset of proteins in similar directions, and the combined knockout produced an amplified response, resulting in potentiation of both upregulated and downregulated protein cohorts. The downregulated cluster was enriched for cell-cycle-associated Hallmarks, suggesting reduced proliferative capacity, whereas the upregulated module was enriched for Glycolysis, ROS Pathway and Apoptosis (Fig. 4B, Supplementary Fig. 3C). Within the upregulated proteins, GPX4, a key glutathione-dependent lipid peroxide reductase^52^ was elevated (Fig. 4C), consistent with increased lipid oxidative pressure. This prompted us to examine proteins annotated to the GO term “Intrinsic Apoptotic Signalling Pathway in Response to Oxidative Stress”^35,36^. These were segregated into two coordinated clusters that distinguished the double knockout from all other genotypes (Fig. 4D). Mitochondrial apoptosis effectors DIABLO and HTRA2, the oxidative-stress-responsive tumour suppressor PML, and SOD2 were upregulated. In contrast, pro-survival factors, including AKT1 were downregulated, further supporting an apoptosis-permissive, oxidative-stress-associated state specific to the double knockout (Fig. 4D). To formally assess genetic interaction effects, we computed per-protein synergy scores across all genotypes. This analysis revealed widespread synergistic behaviour in the double knockout, with 361 downregulated and 314 upregulated proteins exhibiting non-additive responses (Fig. 4E). Synergistically altered proteins were highly enriched for cell-cycle suppression and apoptotic signalling (Supplementary Fig. 3D, E), indicating that combined loss of PHD2 and GSTP1 drives a coordinated, systems-level shift toward growth arrest and cell death that exceeds the effects of either perturbation alone, with potentially lethal consequences for cellular viability.

**Figure 4.**
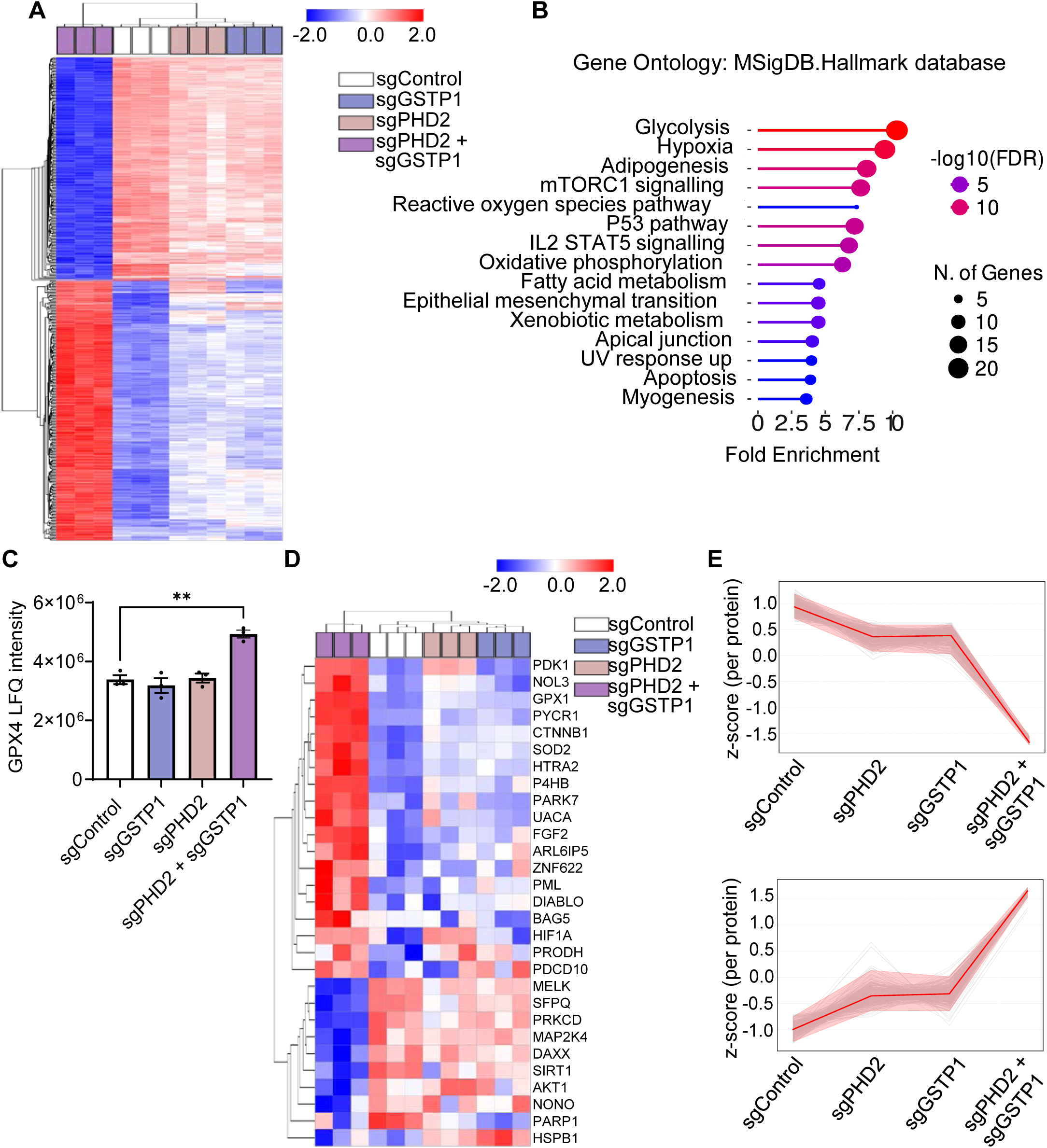
Combined PHD2 And GSTP1 Loss Drives Synergistic Proteomic Changes Associated with Apoptosis. A. Heatmap of top 500 variable proteins in the tested data set. The p-values were calculated using one-way ANOVA. Hierarchical clustering of columns (samples) and rows (proteins detected) was performed using Euclidean distance. B. Protein enrichment analysis of bottom cluster using MSigDB.Hallmark database as a reference. C. MS analyses of GPX4 levels in the cell populations. GPX4 intensity is shown as a label-free quantitation (LFQ) values. Mean + SEM is assessed by unpaired two-tailed Student’s t test, **p<0.01. D. Heatmap of GO term “Intrinsic Apoptotic Signalling Pathway in Response to Oxidative Stress” proteins found in the tested data set. Hierarchical clustering of columns (samples) and rows (proteins detected) was performed using Euclidean distance. E. Synergy profiles of down-regulated (top) and up-regulated (bottom) proteins, expressed as per-protein z-scores. A total of 361 down-regulated (*N* = 361) and 314 up-regulated (*N* = 314) proteins exhibited synergistic behaviour.

Metabolic Imbalance and Apoptotic Vulnerability Following Combined PHD2 And GSTP1 Loss

To assess whether the metabolic signatures inferred from proteomic analyses were reflected at the metabolite level, we performed untargeted metabolomics across all four genotypes (Supplementary Table S4). Unsupervised hierarchical clustering of ANOVA-variable metabolites revealed clear genotype-dependent separation, with the sgPHD2 and sgGSTP1 double knockout forming a distinct metabolic cluster (Fig. 5A). Notably, single knockouts showed partially overlapping profiles, whereas the combined perturbation produced a qualitatively distinct metabolic state, mirroring the non-additive pattern observed at the proteome level. Quantitative analysis of key energetic and redox parameters demonstrated a pronounced imbalance in the double knockout, including an elevated AMP/ATP ratio together with increased GSSG/GSH and altered NAD⁺/NADH and NADP⁺/NADPH ratios (Fig. 5B). These changes were accompanied by coordinated depletion of TCA-cycle intermediates (Fig. 5C), consistent with impaired mitochondrial metabolic capacity and altered redox buffering. Joint enrichment analysis of metabolomics and proteomics data further highlighted altered central carbon and amino-acid metabolism in the double knockout genotype (Supplementary Fig. 4A). Together, these metabolic alterations point to a state of energetic stress and compromised mitochondrial function in the sgPHD2 and sgGSTP1 double knockout, with secondary effects on cellular redox balance. Consistent with this metabolic state, live-cell imaging of cleaved Caspase-3/7 revealed reduced proliferation with a pronounced increase in apoptotic signalling in the double knockout cell line (Fig. 5D, Supplementary Fig. 4B). Notably, this apoptotic response closely resembles the phenotype induced by Molidustat treatment (Fig. 1A, B). Collectively, these data indicate that combined loss of PHD2 and GSTP1 precipitates a metabolic and apoptotic collapse consistent with a synthetic lethal interaction.

**Figure 5.**
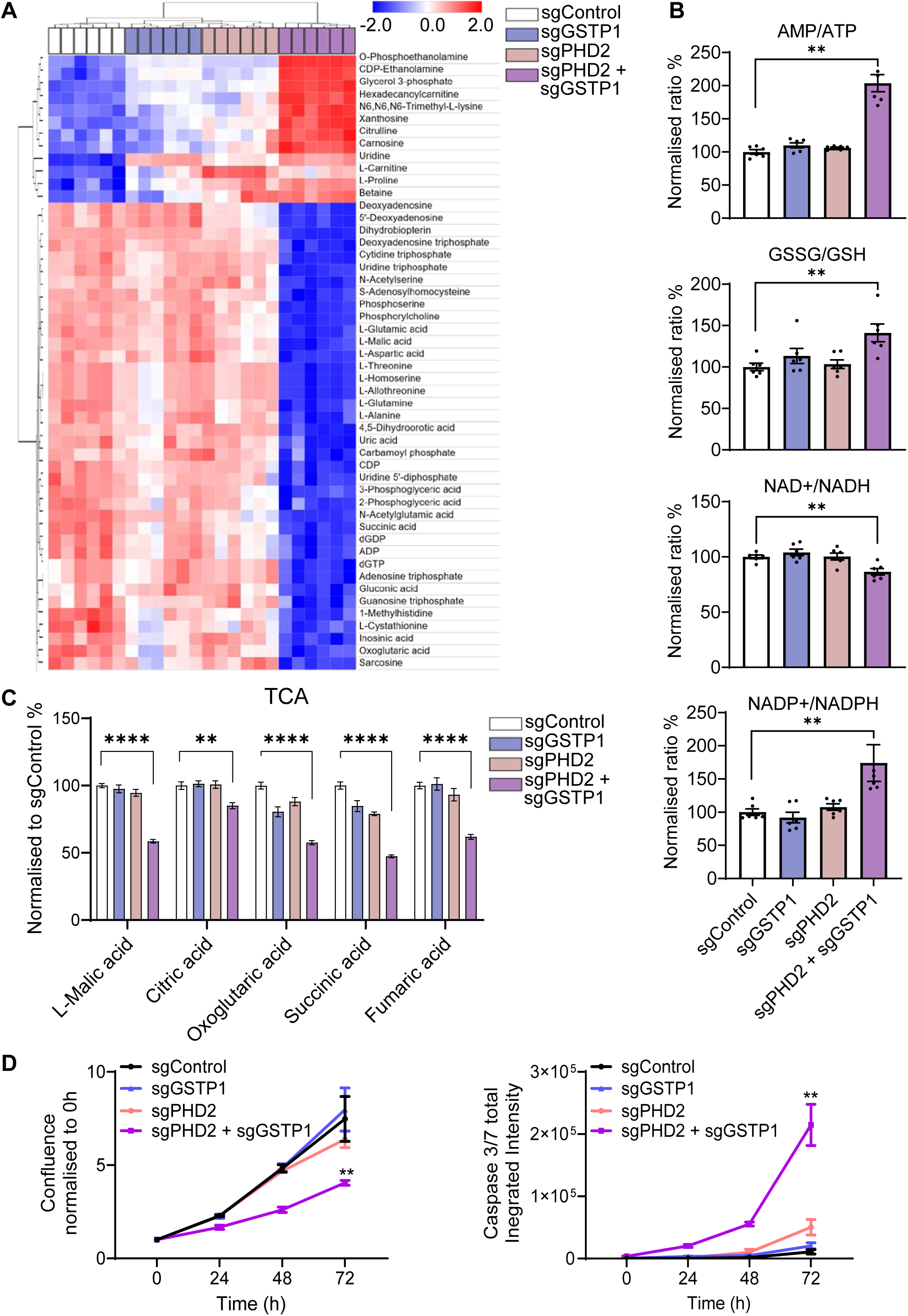
Metabolic Imbalance and Apoptotic Vulnerability Following Combined PHD2 And GSTP1 Loss. A. Heatmap of top 50 variable metabolites in the tested data set. The p-values were calculated using one-way ANOVA. Hierarchical clustering of columns (samples) and rows (metabolites detected) was performed using Euclidean distance. B. Metabolomics analyses of AMP/ATP, GSSH/GSH, NAD+/NADH and NADP/NADH ratios, in the indicated cell lines. Mean + SEM is assessed by unpaired two-tailed Student’s t test, **p<0.01. C. TCA cycle metabolite levels, normalised to sgControl. Mean + SEM is assessed by unpaired two-tailed Student’s t test, **p<0.01, ****p<0.0001. D. Left: Confluence curve normalised to timepoint 0h in the indicated cell lines. Right: Representative Cleaved Caspase-3/7 signal curve in the indicated cell lines. Mean + SEM is assessed by two-way ANOVA, **p<0.01. *N* = 3 biologically independent experiments.

Colonic Organoids Display Genotype-Specific Sensitivity to Molidustat

It has previously been reported that Molidustat treatment reduces tumour growth in an *in vivo* xenograft model of breast cancer^17^. In our CRC *in vivo* model, we detected elevated renal Epo levels, indicating at least partial PHD2 inhibition in the kidney^53^ (Supplementary Fig. 5A). However, Molidustat treatment did not affect survival or tumour characteristics (Supplementary Fig. 5C-F). One potential explanation is insufficient compound delivery to the colon, which we were unable to directly assess.

Given the lack of efficacy in an *in vivo* model, we decided to evaluate the effects of Molidustat in a more relevant system where drug delivery could be better controlled. We therefore employed murine colonic organoids, a stem-cell-derived system that recapitulates epithelial organisation and cellular heterogeneity and enables reliable compound exposure^54^. To determine whether sensitivity to Molidustat was associated with particular colorectal cancer subtypes, we included organoids derived from Genetically Engineered Mouse Models (GEMMs) representing three CMS classifications: Apc^fl/fl^ (CMS2), Apc/Kras/p53 (AKP) (CMS3), Kras/p53/Notch (KPN) (CMS4), in addition to wild-type (WT) controls. Molidustat moderately reduced viability in WT organoids, even at high concentrations, whereas Apc^fl/fl^, AKP and KPN organoids displayed a clear dose-dependent reduction in viability (Fig. 6A). These data suggest that organoids with activated WNT signalling and/or KRAS mutations are sensitive to Molidustat. To assess whether hydroxylase inhibition alone was sufficient to account for this effect, WT and Apc^fl/fl^ organoids were treated with Molidustat or three structurally distinct hydroxylase inhibitors DMOG, JNJ-42041935 and DFO^55,56^. DMOG, a pan-hydroxylase inhibitor, did not affect viability even at 2 mM, whereas JNJ and DFO reduced viability only at high concentrations and did so indiscriminately across genotypes, including WT organoids (Fig. 6B; Supplementary Fig. 6A). In contrast, Molidustat selectively reduced viability in Apc^fl/fl^ organoids (Fig. 6B, Supplementary Fig. 6B), demonstrating that hydroxylase inhibition alone does not recapitulate the selective sensitivity observed with Molidustat. To determine whether Molidustat induced apoptosis in organoids, we assessed Caspase-3/7 activity. Molidustat treated WT organoids showed no detectable activation, whereas approximately half of Apc^fl/fl^ cells displayed increased cleaved Caspase-3/7 signal (Fig. 6C-E), indicating apoptosis as a contributor to reduced viability consistent with activation of a synthetic lethal GSTP1-PHD2 vulnerability in APC-deficient epithelium. To determine whether the stem cell population was affected by Molidustat, we assayed the expression of *Lgr5* and found that mRNA expression was significantly reduced after treatment with the inhibitor, suggesting a reduction in the stem cell population (Fig. 6F). To see whether this reduction of stem cells inhibited the ability regenerate organoids, we conducted a clonogenicity assay from cells derived from vehicle or Molidustat-treated organoids. Consistent with the previous data, Molidustat significantly reduced the clonogenic potential of the organoid-derived cells, suggesting a reduction in the stem cell population (Fig. 6G).

**Figure 6.**
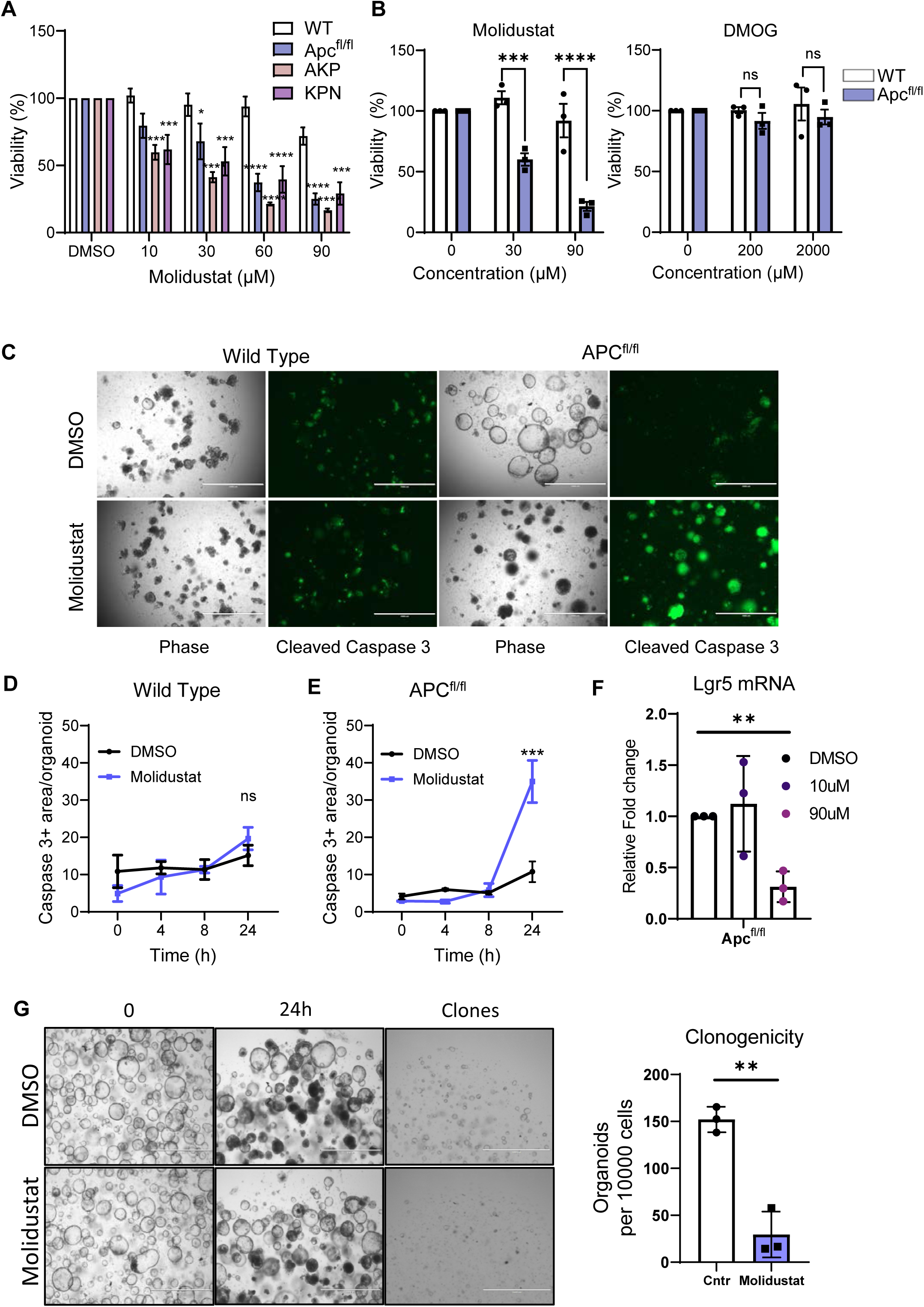
Colonic Organoids Display Genotype-Specific Sensitivity to Molidustat. A. WT, APC^fl/fl^(CMS2), AKP (CMS3) and KPN (CMS4) organoids were treated with increasing concentrations of Molidustat for 48 h. Viability percentage was assessed using an MTT Assay. Mean + SEM is assessed by Two-way Anova with Dunnett’s Correction for Multiple Comparisons, *p<0.05, ***p<0.001, ****p<0.0001. B. 2-day old wild type (WT) or APC^fl/fl^ organoids were treated for 48h hours with the indicated PHD inhibitors. Organoid viability percentage was assessed with an MTT assay. Mean + SEM is assessed by unpaired two tailed Student’s t-test, ***p<0.001, ****p<0.0001, (ns) non-significant. C. Representative images of Cleaved Caspase-3/7 signal in wild type and APC^fl/fl^ organoids treated with DMSO or Molidustat at 24 h timepoint. Scale bar: 1000 μm. D-E: Quantification of Cleaved Caspase-3/7 signal area in organoids at the indicated timepoints. Mean + SEM is assessed by unpaired two tailed Student’s t-test, ***p<0.001, (ns) non-significant. *N* = 3 biologically independent experiments. F. Normalised RT-qPCR quantification of Lgr5 mRNA expressed in APC^fl/fl^ organoids treated with DMSO or Molidustat at 24 h timepoint two tailed Student’s t-test, **p<0.01. *N* = 3 .G. Representative images of APC^fl/fl^ organoids treated with DMSO or Molidustat at two timepoints and clones grown from organoids treated for 24 h with DMSO or Molidustat. Quantification of clones in on the right, two-tailed Student’s t-test, **p<0.01, (ns). *N* = 3

Overall, these findings demonstrate that Molidustat selectively reduces viability in colonic organoids with distinct mutational backgrounds, particularly those with activated WNT signalling and/or KRAS pathway alterations, and that this effect is not reproduced by hydroxylase inhibition alone.

## Discussion

In this study, we identify GSTP1 as a previously unrecognised off-target interactor of Molidustat and reveal a synthetic lethal interaction between GSTP1 and PHD2 whose combined inhibition is selectively deleterious in APC-mutant colorectal cancer cells. Using TPP, chemical proteomics, CRISPR knockouts, and multi-omics profiling, we show that simultaneous inhibition of GSTP1 and PHD2 disrupts cellular homeostasis and induces oxidative-stress and apoptosis, closely phenocopying the response to Molidustat treatment. Notably, GSTP1 and PHD2 loss each affect overlapping yet mechanistically distinct proteomic networks, providing a plausible basis for the emergent synthetic interaction. Synthetic lethal relationships of this type, where two enzymes regulate convergent pathways through independent mechanisms, are well documented, exemplified by BRCA1 and PARP in DNA-damage repair^57,58^.

Mechanistically, our proteomic and metabolomic analyses indicate that the loss of dual GSTP1/PHD2 leads to ROS accumulation, elevated GSSG/GSH ratios, induction of oxidative-stress-associated apoptotic mediators, and downregulation of pro-survival pathways. Together, these coordinated alterations define a potential redox-collapse mechanism that underpins this synthetic interaction and explains the apoptotic phenotype observed upon both genetic double knockout and Molidustat exposure. However, the magnitude of oxidative stress suggests that additional oxidative stress-associated cell death mechanisms may also contribute. While genetic rescue experiments would provide definitive confirmation of on-target specificity, the pronounced loss-of-fitness and apoptotic phenotype observed upon combined PHD2 and GSTP1 loss limited the feasibility of establishing stable rescued double-knockout populations, and therefore represents a limitation of the current study. Prior reports show that the pan-PHD inhibitor DMOG perturbs mitochondrial function before inducing HIF targets^14^, raising the possibility that GSTP1 may buffer mitochondrial stress in this context. Notably, hypoxia and HIF-targeting agents have recently been recognised as potential GSTP1 inhibitors based on molecular docking analyses, suggesting that GSTP1 engagement may represent a broader, previously underappreciated feature of compounds developed to modulate hypoxia signalling^59^.

Although synthetic lethality is well established across cancer types, including examples such as PARP-BRCA, ATR-Wee1, and WRN in microsatellite-instable cancers ^57,60,61^, the GSTP1-PHD2 interaction has not been previously described. This expands the spectrum of actionable dependencies in APC-mutant CRC. To fully validate this axis *in vivo*, two considerations are important. First, Molidustat delivery to the colon in our CRC models was suboptimal, but alternative strategies, such as implantable microdevices previously used to deliver PHD inhibitors directly into tumour xenografts^62^ provide feasible routes to achieve effective local exposure. Second, synthetic lethal relationships identified *in vitro* do not always translate to animal models^63^, underscoring the need for an APC-deficient CRC model with dual GSTP1/PHD2 loss. Together, these approaches will enable a rigorous assessment of the GSTP1-PHD2 dependency *in vivo* and strengthen the translational relevance of this vulnerability.

The clinical characteristics of Molidustat strengthen its translational potential. Long-term studies demonstrate good tolerability for up to 36 months in chronic kidney disease^64^, an appealing safety profile given the toxicity of many frontline CRC treatments ^65,66^. Beyond CRC, Molidustat also exerts anti-tumour effects in breast cancer, where it reduces cell viability, induces cell-cycle arrest, enhances the activity of chemotherapeutic agents, and suppresses tumour growth in xenografts^17^. Notably, Molidustat robustly stabilises HIF-1α and induces VEGF to levels comparable to hypoxia, presenting an opportunity for combination strategies that counterbalance HIF-driven angiogenesis. Bevacizumab, which blocks VEGF-mediated angiogenic signalling^67^, represents one such rational partner. Molidustat also enhances the efficacy of gemcitabine, an antimetabolite that inhibits DNA synthesis, supporting exploration of this combination in additional cancer contexts^17,68^. Furthermore, identification of PHD2 as a druggable dependency in KRAS-driven lung cancer ^69^ suggests potential synergy with KRAS inhibitors, including in APC/KRAS co-mutant CRC. By simultaneously exploiting the GSTP1-PHD2 synthetic lethal axis and modulating hypoxia signalling, Molidustat offers a mechanistically grounded entry point for multi-agent regimens that incorporate VEGF-pathway inhibitors, chemotherapeutics, or KRAS-targeted therapies.

A further consideration raised by our findings concerns the isoform selectivity of Molidustat. Although developed as a PHD2 inhibitor, Molidustat retains appreciable activity against PHD1 and PHD3^70^, and the relative contributions of each isoform in the gut are non-redundant and, in some contexts, opposed. Genetic deletion of PHD1, but not PHD2 or PHD3, protects mice from DSS-induced colitis by limiting epithelial apoptosis and preserving barrier function, and PHD1 protein is elevated in mucosal biopsies from patients with ulcerative colitis and Crohn’s disease^71^. In colitis-associated cancer, PHD1 loss restrains tumour growth, whereas PHD2 haploinsufficiency has a complex phenotype resulting in increased proliferation, cell death, and modifies the immune response^72^. Notably, this pattern of isoform engagement is shared with other pan-PHD inhibitors that did not phenocopy Molidustat in our screens, indicating that PHD isoform profile alone is insufficient to explain its distinctive activity and pointing instead to the GSTP1 off-target engagement as the key distinguishing feature. Nonetheless, the compartment-specific roles of PHD1 and PHD2 argue for localised colonic delivery strategies, such as the microdevice-based approaches discussed above, which would concentrate Molidustat at the APC-mutant epithelium to exploit the GSTP1–PHD2 synthetic lethal axis while limiting systemic exposure and the potentially deleterious consequences of systemic PHD2 inhibition.

Several GSTP1 inhibitors are currently in preclinical development, and our findings suggest that their antineoplastic activity may be potentiated through concurrent disruption of PHD2. Thus, a dual-inhibition strategy targeting both GSTP1 and PHD2 may yield synergistic efficacy, particularly in genetic backgrounds that rely on coordinated redox and hypoxia-adaptation programmes. In this context, Molidustat, despite its originally intended selectivity toward PHD2, emerges as a versatile candidate for rational combination approaches, not only due to its clinical tractability but also because of its unexpected engagement of GSTP1. Collectively, these observations broaden the therapeutic landscape for exploiting oxidative-stress vulnerabilities across multiple cancer subtypes.

## Author Contributions

Conceptualization, K.B.M and A.V.K. Methodology and Investigation C.A., A.N.M., J.M.J., A.B.A., J.W., A.Y., A.P.L, C.J, A.U.B., K.B.M and A.V.K. Writing - original draft, A.N.M. and A.V.K. Writing - review and editing, all authors. Funding acquisition, K.B.M, A.V.K.

The University of Edinburgh wishes to acknowledge the validity of Pelago’s CETSA patent (EP 2 699 910 B1) entitled “Methods For Determining Ligand Binding To A Target Protein Using A Thermal Shift Assay” (the “Patent”) and the use of the Cellular Thermal Shift Assay (CETSA) process described in the Patent in this manuscript.

## Acknowledgments

All illustrations were created with BioRender.com. We thank Prof Owen Sansom (Cancer Research UK Scotland Institute) and Dr Ömer H Yilmaz (Massachusetts Institute of Technology) for providing organoid models. We are grateful to members of the Myant and von Kriegsheim laboratory for their discussions and critical reading of the manuscript. C.A was supported by a CRUK studentship. K.B.M was supported by Cancer Research UK (CRUK) under a Career Development Fellowship (A19166 to K.B.M.) and a Small Molecule Drug Discovery Project Award (A25808 to K.B.M.) and a European Research Council under Starting Grant (COLGENES–715782 to K.B.M.). A.V.K. acknowledged MRC Equipment grant (MR/X01293X/1), BBSRC Alert (BB/X019160/1) and Welcome Trust Multi-user Equipment Grant (grant ID: 208402/Z/17/Z) for funding the instrumentation.

