## Supplementary Figures for "Molidustat Targets a Synthetic Lethal Vulnerability in APC-Mutant Colorectal Cancer through GSTP1 and PHD2 Co-Inhibition"

### Molidustat Induces Cell Death in APC-Mutant Colorectal Cancer through GSTP1 and PHD2 Co-Inhibition

### **Table of contents**:

Figure S1 - Molidustat induces HIF/signalling and increases PHD2 thermal stability

Figure S2 - Molidustat Inhibits GSTP1 Activity

Figure S3 - Proteomics analyses of HT29 knockout cell models

Figure S4 - Metabolomics analyses of HT29 knockout cell models

Figure S5 - Evaluation of Molidustat in CRC *in vivo* model

Figure S6 - The effects of different PHD inhibitors on murine colonic organoids viability

Table S1 - Hits from TPP screen (Easy50 method) (XLSX)

Table S2 - IOX4 screen Mass Spectrometry proteomics FragPipe Search Output and analysis files (XLSX)

Table S3 - HT29 Mass Spectrometry proteomics DIA-NN Search Output and analysis files (XLSX)

Table S4 - HT29 Mass Spectrometry metabolomics Search Output and analysis files (XLSX)

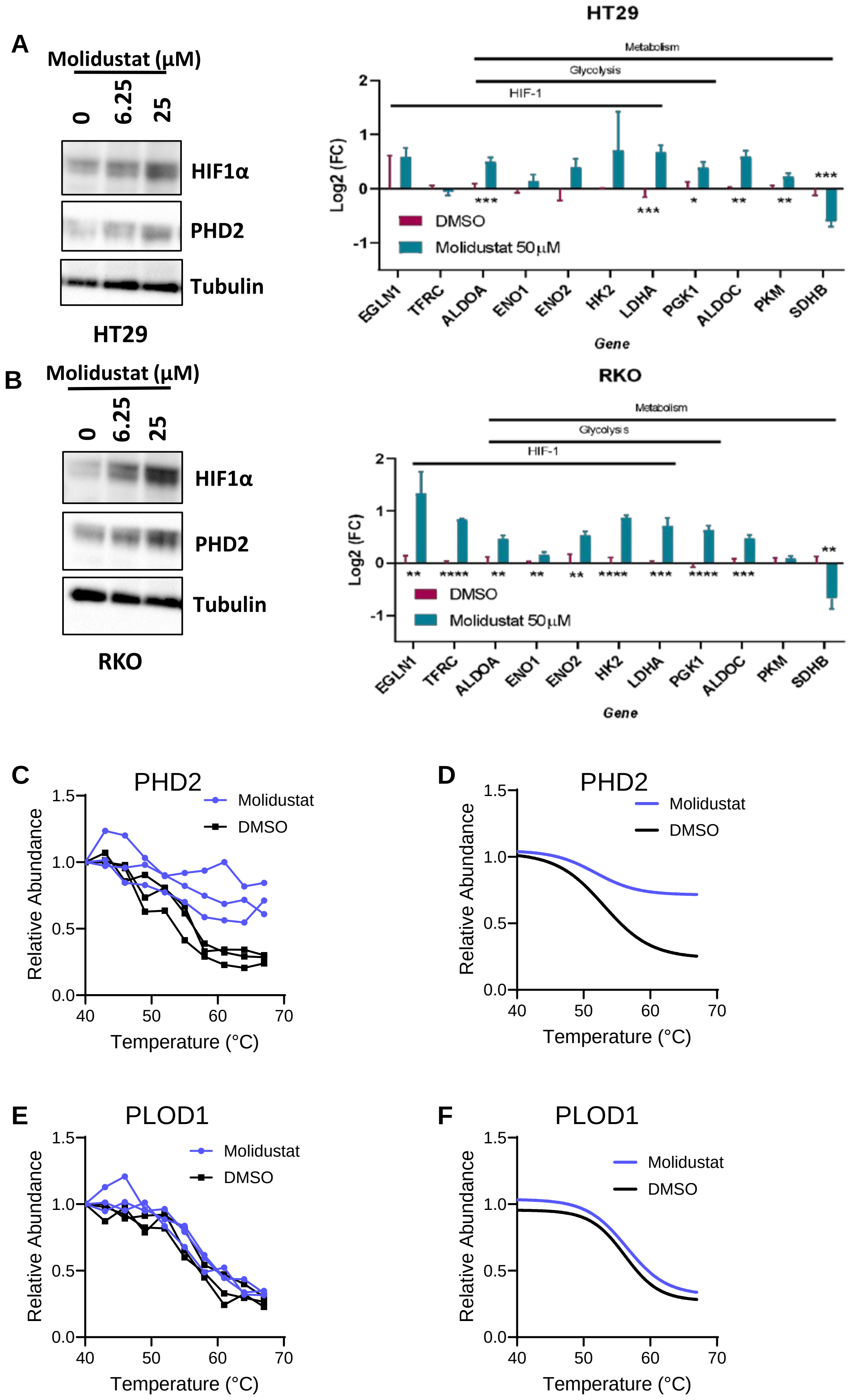


**Figure S1. Molidustat induces HIF/signalling and increases PHD2 thermal stability.** A&B. Western blots and Protein expression changes investigating HIF1α and PHD2 levels with increasing concentrations of Molidustat in HT29 and RKO cells 16 h B. Protein levels of HIF-responsive genes upon treatment of Molidustat, 16 h C-D Raw values and sigmoidal curve fits for PHD2 in samples treated with Molidustat (purple) or DMSO (blue). E-F Raw values and sigmoidal curves for PLOD1 in samples treated with Molidustat (purple) or DMSO (blue). *N* = 3 biologically independent experiments.

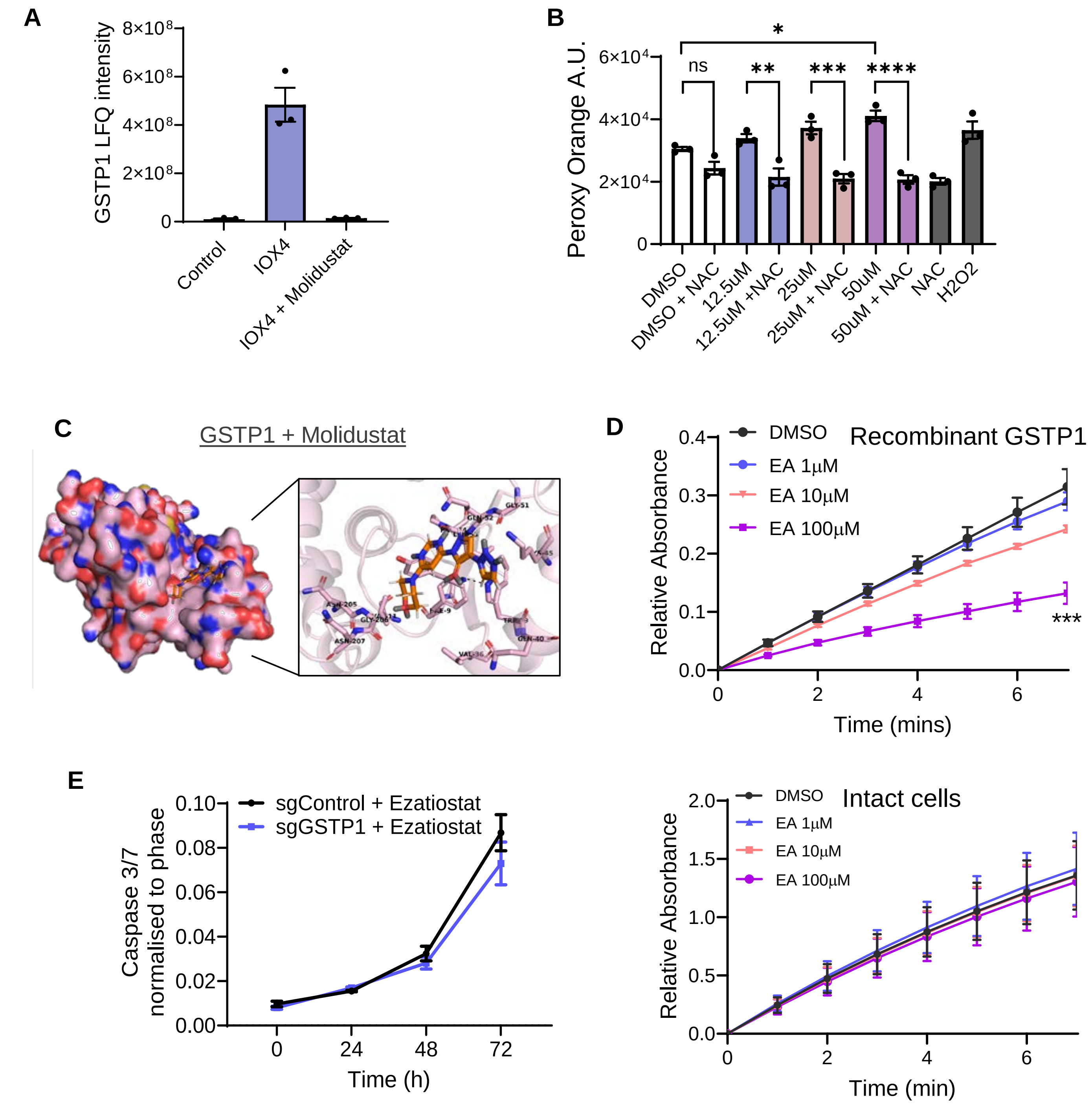


**Figure S2. Molidustat Inhibits GSTP1 Activity.** A. MS analyses of GSTP1 levels in the indicated conditions. Protein intensity is shown as a label-free quantitation (LFQ) values. B. HT29 cells were treated with increasing concentrations of Molidustat. ROS were measured using Peroxy Orange 1 (PO1) and expressed as arbitrary units (AU). At 50 µM Molidustat, a significant increase in PO1 fluorescence was observed. Co-treatment with 5 mM N-acetylcysteine (NAC), a ROS scavenger, abolished the Molidustat-induced signal. Mean + SEM is assessed by one-way ANOVA, *p<0.05, **p<0.01, ***p<0.001 , ****p<0.0001, (ns) non-significant. *N* = 3 biologically independent experiments. C. Molecular docking of Molidustat to GSTP1 protein. Residues in the active site are highlighted on the right. D. Recombinant (left) or cell-based (right) GSTP1 assay with ethacrynic acid (EA) or DMSO; time-course of CDNB-GSH conjugation monitored at 340 nm and plotted as relative absorbance (arbitrary units, AU). E. Cleaved Caspase-3/7 signal curve in the indicated cell lines. Mean + SEM is assessed by two-way ANOVA, *N* = 1 biologically independent experiment.

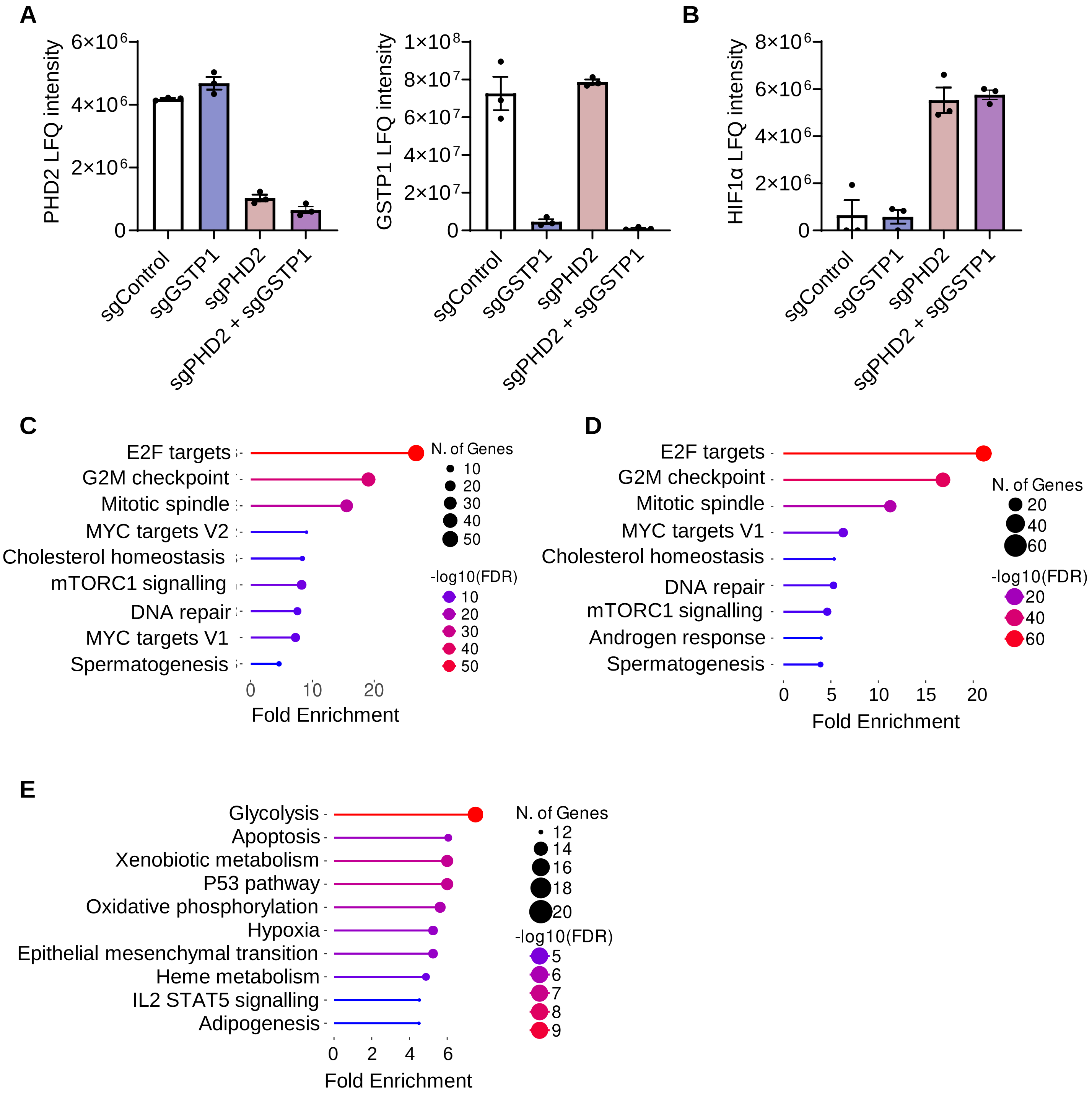


**Figure S3. Proteomics analyses of HT29 knockout cell models.** A. MS analyses of PHD2 (left), or GSTP1 (right) levels in the cell populations. Protein intensity is shown as a label-free quantitation (LFQ) values. B MS analyses of HIF1A levels in the cell populations. HIF1A intensity is shown as a label-free quantitation (LFQ) values. C. Protein enrichment analysis of top cluster from Figure 4A using MSigDB.Hallmark database as a reference. D. Protein enrichment analysis of synergistic, down-regulated proteins (as seen in Fig. 3E) using MSigDB.Hallmark database as a reference. D. Protein enrichment analysis of synergistic, up-regulated proteins (as seen in Fig. 3E) using MSigDB.Hallmark database as a reference.

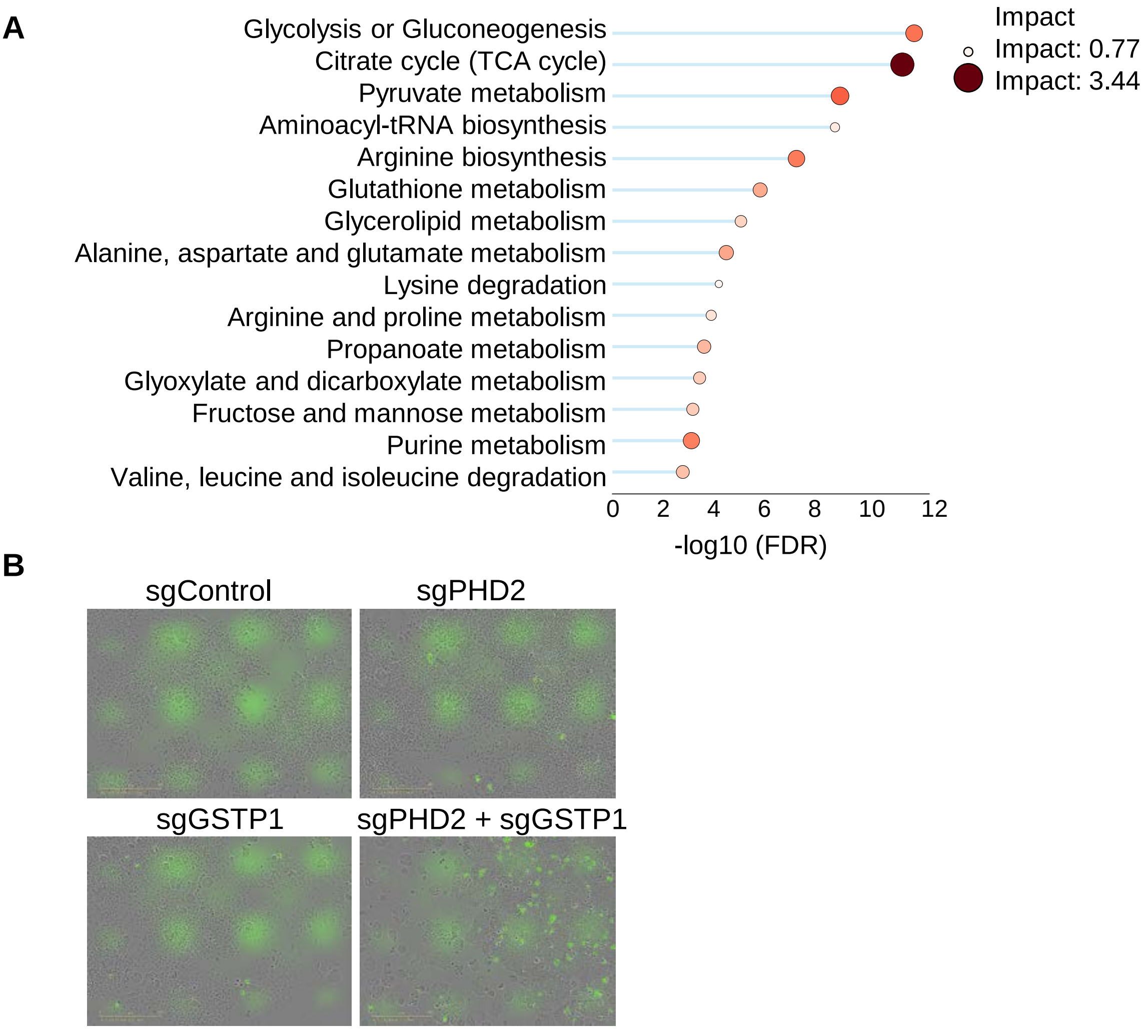


**Figure S4. Metabolomics analyses of HT29 knockout cell models.** A. Enrichment analysis of metabolomics and proteomics data, showing top 15 enriched metabolic processes in sgPHD2 and sgGSTP1 cell line comparing to sgControl. B. Representative merged images of Phase and Cleaved Caspase 3/7 signal in the indicated cell lines. Scale bar: 400 μm. *N* = 3 biologically independent experiments.

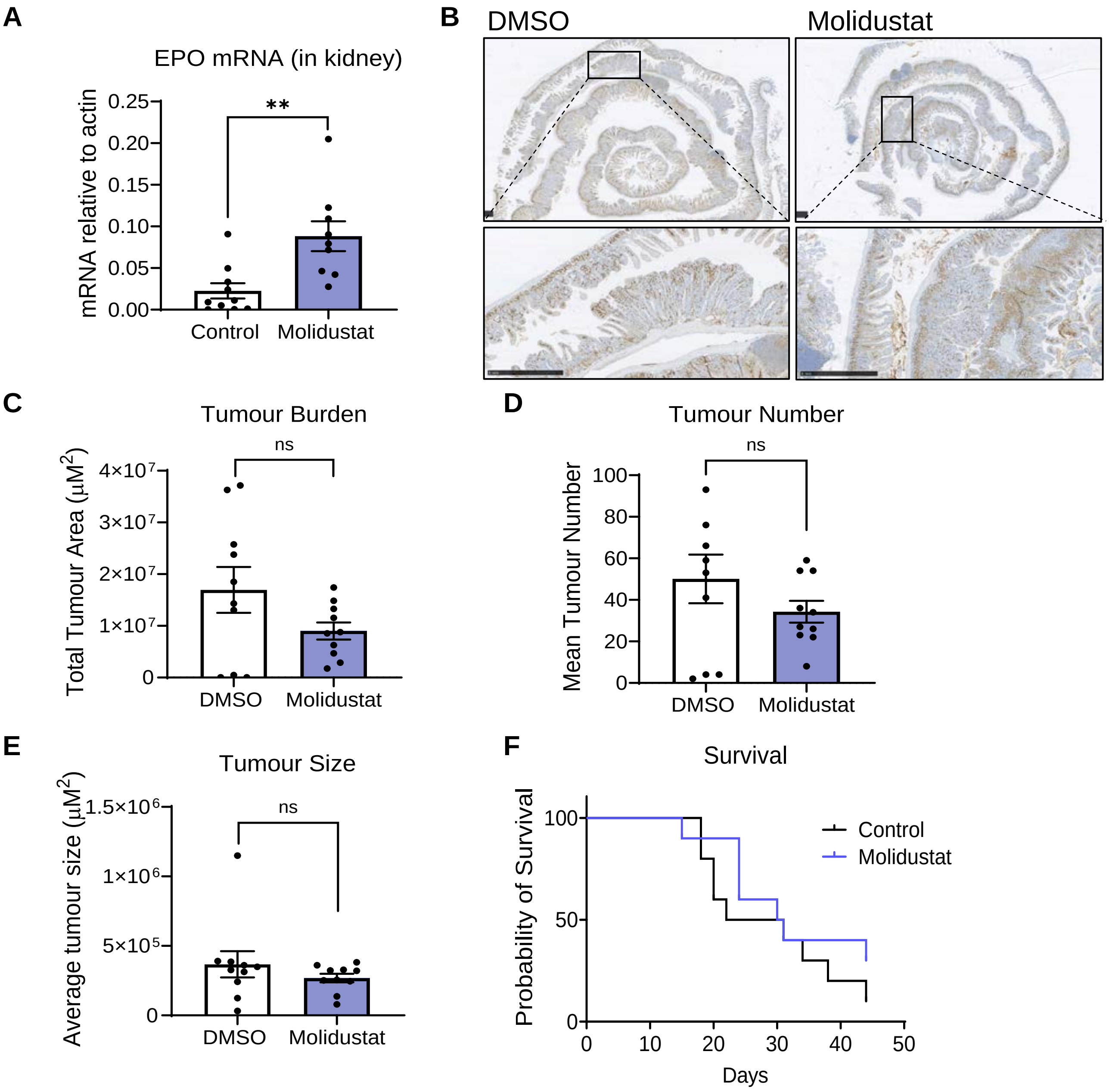


**Figure S5. Evaluation of Molidustat in CRC *in vivo* model.** A. Epo mRNA levels after Molidustat treatment. Mean + SEM is assessed by Mann-Whitney U test, **p<0.01. B. Representative immunohistochemistry images of rolls of the upper section of the intestinal epithelium with a large number of macroscopic and microscopic tumours. Sections were stained with BrdU to stain DNA of actively dividing cells. C. Tumour burden of mice treated with DMSO or Molidustat. Mean + SEM is assessed by unpaired two tailed Student's t-test, (ns) non-significant. D. Tumour number in mice treated with DMSO or Molidustat. Mean + SEM is assessed by unpaired two tailed Student's t-test, (ns) non-significant. E. Tumour size of mice treated with DMSO or Molidustat. Mean + SEM is assessed by unpaired two tailed Student's t-test, (ns) non-significant. F. Survival curve of mice treated with DMSO or Molidustat. Log-rank Mantel Cox test, p=0.2924. *N* = 10 DMSO, *N* = 10 Molidustat

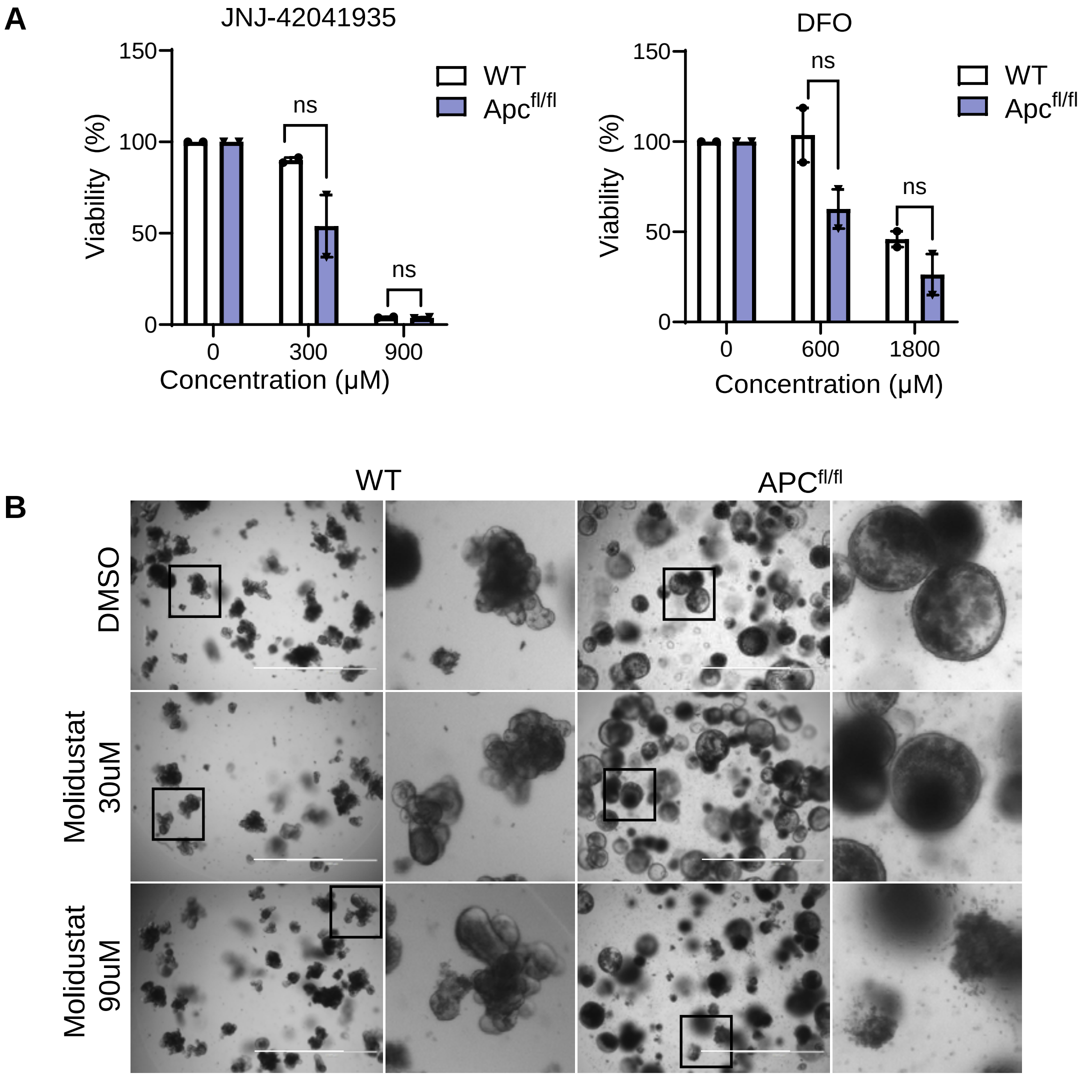


**Figure S6. The effects of different PHD inhibitors on murine colonic organoids viability.** Wild type (WT) or APC organoids viability after 48 hour treatment with the indicated PHD inhibitors. Organoid viability percentage assessed using MTT assay. Mean + SEM is assessed by two-way ANOVA, (ns) non-significant. *N* = 2 biologically independent experiments. B. Representative images of WT and APC organoids treated with 30 and 90μM Molidustat. Scale bar: 1000 μm. *N* = 3 biologically independent experiments.
